# Comparison of gene clustering criteria reveals intrinsic uncertainty in pangenome analyses

**DOI:** 10.1101/2022.09.25.509376

**Authors:** Saioa Manzano-Morales, Yang Liu, Sara González-Bodí, Jaime Huerta-Cepas, Jaime Iranzo

## Abstract

**Background:** A key step for comparative genomics is to group open reading frames into functionally and evolutionarily meaningful gene clusters. Gene clustering is complicated by intraspecific duplications and horizontal gene transfers, that are frequent in prokaryotes. In consequence, gene clustering methods must deal with a trade-off between identifying vertically transmitted representatives of multi-copy gene families (recognizable by synteny conservation) and retrieving complete sets of species-level orthologs. We studied the implications of adopting homology, orthology, or synteny conservation as formal criteria for gene clustering by performing comparative analyses of 125 prokaryotic pangenomes.

**Results:** Clustering criteria affect pangenome functional characterization, core genome inference, and reconstruction of ancestral gene content to different extents. Species-wise estimates of pangenome and core genome sizes change by the same factor when using different clustering criteria, which allows for robust cross-species comparisons regardless of the clustering criterion. However, cross-species comparisons of genome plasticity and functional profiles are substantially affected by inconsistencies among clustering criteria. Such inconsistencies are driven not only by mobile genetic elements, but also by genes involved in defense, secondary metabolism, and other accessory functions. In some pangenome features, the variability attributed to methodological inconsistencies can even exceed the effect sizes of ecological and phylogenetic variables.

**Conclusions:** Choosing an appropriate criterion for gene clustering is critical to conduct unbiased pangenome analyses. We provide practical guidelines to choose the right method depending on the research goals and the quality of genome assemblies, and a benchmarking dataset to assess the robustness and reproducibility of future comparative studies.

## Background

Recent advances in sequencing have revolutionized the study of microbial ecology and evolution by providing access to thousands of high-quality genomic sequences. A major consequence of the surge in microbial genomic data has been the introduction of the concept of pangenome, that is, the set of all genes found in the genomes of a taxonomic group [1,2]. Pangenomes have quickly gained importance in eco-evolutionary research because they provide valuable information about the functional capabilities accessible to the strains of the same species and their propensity to gain and lose genes [3] Therefore, the study of pangenomes can shed light on the interactions among genome plasticity, niche diversity, and adaptability to environmental changes [4–7]. Moreover, identifying the set of genes present in all the genomes of a species (the core genes) has proved helpful to define core functions and build high-resolution species trees [8–10].

To study the pangenome of a species, researchers first collect all available high-quality genomes and run gene prediction tools to identify open reading frames (ORF). The ORF are then clustered into groups of “equivalent” genes, that serve as the basis for subsequent cross-genome comparisons. To obtain meaningful results, it is critical that gene clusters represent coherent units, both from functional and evolutionary perspectives. The simplest conceptual approach to gene clustering is based on homology. Two sequences are homologous if they derive from a common ancestral sequence. Beyond this qualitative definition, different types of homology can be established based on the evolutionary history of each gene. Because most gene families have experienced duplications throughout evolution, it is usual to find homologous genes with multiple representatives per genome, termed paralogs. Paralogs often display functional divergence and accelerated evolutionary rates [11,12]. As a result, homology alone does not guarantee functional and evolutionary homogeneity of gene clusters. Instead, for most purposes, it is desirable to subdivide homologs into higher-resolution and more homogeneous clusters encompassing orthologous sequences [13,14]. Orthologs are genes that share a single common ancestor at the time of speciation [13,15]. Accordingly, the orthology criterion discriminates paralogs that duplicated before the last speciation event (Fig. 1). Because of the connection between orthology and speciation, orthology is the most natural grouping criterion for cross-strain comparative genomics and phylogenomic analyses. In practice, it is challenging to apply a strict orthology criterion to pangenome studies due to the lack of accurate reference species trees and the high computational burden of building thousands of single-gene trees, one for each group of homologs. To circumvent these difficulties, heuristic algorithms and reference databases of orthologous genes have been developed over the last decade [16–18]. Together with homology and orthology, a third criterion to further refine equivalence among genes is synteny conservation. According to the synteny criterion, two genes (typically orthologs) are grouped together if they share the same gene neighborhood in different genomes. Notably, synteny conservation can help discriminate between vertically and horizontally transmitted copies of a gene, or between multiple orthologs resulting from within-species gene expansions (Fig. 1).

**Figure 1:**
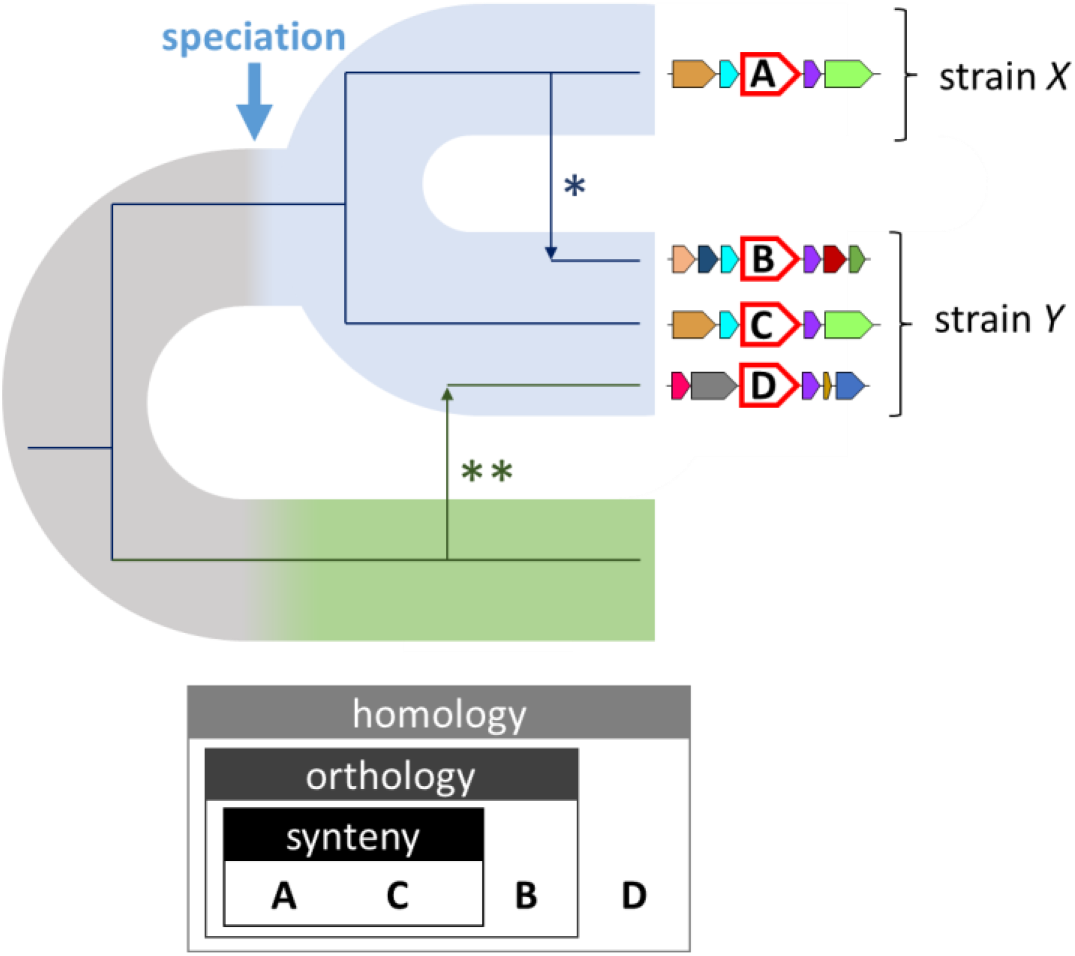
Homology, species-level orthology, and synteny conservation. The phylogeny of a gene family that has experienced intra-species HGT (*) and inter-species HGT (**) is superimposed on a species tree (colored ribbons). In the species tree, speciation is followed by diversification into two strains (blue), and a sister species is represented in green. Gene neighborhoods are displayed at the leaves of the gene tree, with the gene of interest in the middle (labeled A-D, larger size, and red border). All sequences (A-D) are homologs. However, only A, B, and C are species-level orthologs, because they descend from the same ancestral sequence at the time of speciation. Of those, A and C were vertically transmitted and share conserved gene neighborhoods (synteny). Sequences B and C are in-paralogs, whereas C and D are out-paralogs.

It is widely accepted that the quantitative results of single-species pangenome analyses depend on the method (and parameters) used to build the pangenome. Such dependency can be traced back to the strategy adopted to identify gene clusters. As intuitively expected, the size of the pangenome (that is, the number of clusters) increases and the size of the core genome decreases as the sequence similarity threshold used to define homology becomes more and more stringent [19–21]. However, comparative studies implicitly assume that qualitative trends and order relationships are robust with respect to the choice of a particular criterion. The existence of such qualitative robustness merits a critical evaluation, given the deep conceptual differences among homology, orthology, and synteny conservation, the heterogeneity of evolutionary rates across genes and species, and the high rates of horizontal gene transfer (HGT) observed in some genomes [22–24].

The possibility that homology, orthology, and synteny-based gene clustering produce incongruent results in comparative pangenome studies is not just a technical caveat. Instead, intraspecific HGT and gene duplications raise the fundamental question of what equivalence class (homology, orthology, synteny, or something else) best captures the essentially dynamic nature of pangenome evolution [25]. This conceptual conundrum is especially clear in what concerns the splitting of multicopy orthologs based on their gene neighborhoods. Let us consider, for example, a gene that has experienced recent duplications or intraspecific HGT. For what purposes should all copies be included in the same group, following the standard orthology criterion? For what other purposes should the duplicated or horizontally transferred copy be assigned to a new cluster, based on synteny considerations? The orthology criterion appears better to assess pangenome diversity, for which it is not desirable to count a recently duplicated gene twice [26], or to study gene duplication and intraspecific HGT, which first requires classifying multicopy genes as members of the same group [27]. In contrast, the synteny criterion would be more appropriate to select marker genes for phylogenomic analyses or identify vertically transmitted members of mobile gene families.

In this study, we show that the choice of a particular clustering criterion can have notable consequences on downstream comparative analyses. To stress the fact that the optimal criterion is somewhat arbitrary and dependent on the main research goal, we adopt the method-agnostic term *Operational Gene Cluster* (OGC) as an umbrella that includes homologs, classical orthologs (possibly inferred through different methods), and vertically transmitted subsets of orthologs with conserved synteny. Depending on the particular choice, OGC may imply different degrees of functional equivalence and shared ancestry.

State-of-the-art methods for OGC construction implement different strategies to deal with sensitivity-vs-specificity trade-offs at manageable computational cost. The simplest approaches (e.g., CD-HIT [28] and MMseqs2 [29]), build groups of homologous genes by applying a pre-defined similarity threshold to the amino acid sequences encoded by a set of ORF. More sophisticated tools include additional steps to discriminate true orthology from other ways of homology. Such discrimination can be attained through two major strategies: (a) by building gene-level phylogenetic trees or sequence similarity networks, on which some heuristic rules are applied to resolve subsets of genes with shared ancestry [30]; and (b) by subclustering homologous groups based on their gene neighborhoods, under the assumption that synteny is locally conserved at the evolutionary timescales that are relevant to within-species diversification. By design, the former approach, which is adopted by reference databases like COG [16] and eggNOG [18], the orthology detection tool OrthoFinder [31], and the pangenome analysis suite panX [32], is better suited to assess true orthology in gene families affected by intraspecific duplications and HGT. However, its high computational burden makes it inefficient to deal with large genomic datasets. On the other side, synteny-based approaches (such as the one implemented by the popular pangenome analysis tools Roary [33] and PanOCT [34]) are faster and less resource intensive, although their results may deviate from the classical concept of orthology (by missing true orthologs) when applied to highly dynamical genomes or regions of the genome with poor synteny conservation. Phylogeny-aware and synteny-based methods are not mutually exclusive [35], although they are rarely performed together due to computational constraints. A summary of these and other popular tools for OGC construction and their application to pangenome analysis is provided in Suppl. Table S1.

Despite their distinct conceptual underpinnings, different types of OGC are sometimes just viewed as exchangeable heuristic approaches that approximate the concept of orthology in a computationally tractable manner. We tested to what extent such an assumption is true and found that some properties of the pangenome, mostly concerning its size and the identity of the core genome, are indeed robust. However, pangenome properties that are related to its fluidity (that is, the genomic variability among strains) can be greatly affected, leading to relatively poor correlation in the results of comparative genomic analyses conducted with different methods. Interestingly, this does not only affect mobile genetic elements, but also genes involved in non-selfish functions.

## Results

### Method-dependent variation and intrinsic uncertainty in pangenome size and diversity

We investigated 5 major methods for *de novo* species-wise OGC construction that qualitatively differ in the strategies for gene clustering and paralog discrimination (Table 1). All *de novo* methods start from a collection of open reading frames (ORF) and cluster them according to a predefined identity threshold that can be set to include more or less distant homologs. The resulting clusters are then processed to split paralogs into separate OGC. The methods included in this study represent three alternative approaches implemented by some of the most popular tools for pangenome analysis: orthology-based clustering (implemented by panX and OrthoFinder), synteny-based clustering (implemented by Roary), and homology-based clustering (implemented by CD-HIT and MMseqs2, which is the OGC construction module used by PanACoTA [36] and PPanGGOLiN [37]. In the case of Roary, CD-HIT, and MMseqs2, we also explored the effect of introducing two different identity thresholds for the initial clustering step (by design, panX and OrthoFinder do not filter gene clusters based on sequence identity but on their e-value). The optional settings used with each tool are detailed in the methods section and summarized in Suppl. Table S2.

**Table 1:**
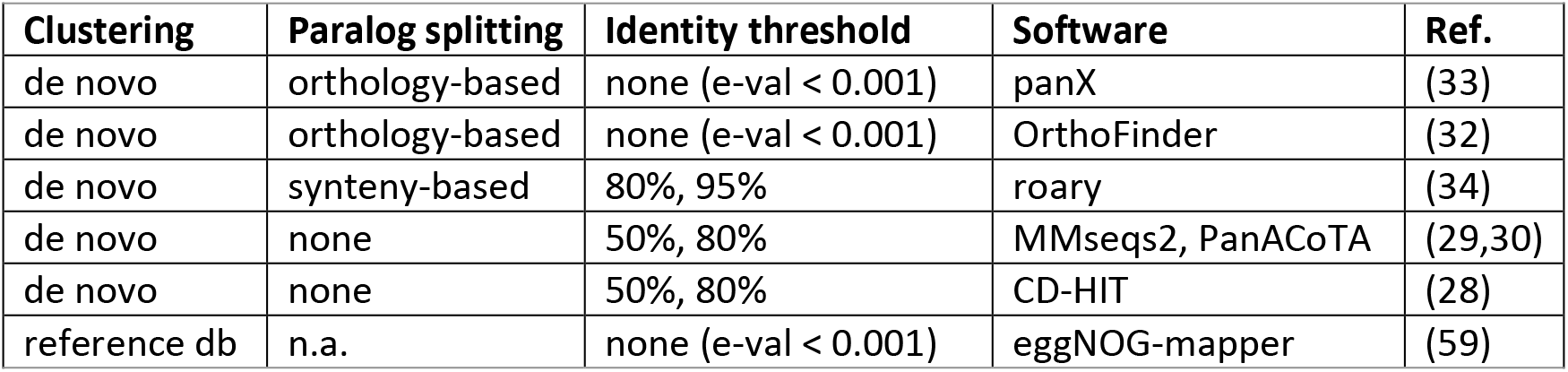
OGC generation strategies and tools used in this study.

| Clustering | Paralog splitting | Identity threshold | Software | Ref. |
| --- | --- | --- | --- | --- |
| de novo | orthology-based | none (e-val < 0.001) | panX | (33) |
| de novo | orthology-based | none (e-val < 0.001) | OrthoFinder | (32) |
| de novo | synteny-based | 80%, 95% | roary | (34) |
| de novo | none | 50%, 80% | MMseqs2, PanACoTA | (29,30) |
| de novo | none | 50%, 80% | CD-HIT | (28) |
| reference db | n.a. | none (e-val < 0.001) | eggNOG-mapper | (59) |

In addition, we considered a fast orthology prediction method (eggNOG-mapper) that maps ORF to a reference database of orthologous groups. This reference-based method was not originally intended for pangenome analysis and has some important limitations. First, because the eggNOG database was built using only one representative genome per species, it does not account for variability across strains. Second, because orthologous groups in the eggNOG database were required to contain sequences from at least 3 species, mapping ORF to eggNOG automatically excludes sequences without known homologs and sets a hard limit for OGC taxonomic resolution at the genus level. Still, reference-based orthology assignments are nowadays highly efficient and scalable to large (meta)genomic datasets. Therefore, we included eggNOG-mapper in this study to assess its performance compared to *sensu stricto* pangenome reconstruction methods.

By applying these methods, we obtained 9 alternative sets of species-wise OGC for 124 bacterial and 1 archaeal species (DOI:10.5281/zenodo.7387758). These species, defined according to the phylogenetically consistent classification scheme established by the Genome Taxonomy Database (GTDB) [38], were selected to cover every genus in the GTDB, with the condition that there were at least 15 high-quality genomes available per species. We quantified the discordance between the OGC produced by each pair of methods by means of the normalized variation of information (NVI), a measure that takes values between 0, if the two sets of OGC display a one-to-one correspondence, and 1, if they are completely independent. The hierarchical clustering of methods based on this measure reveals that the discordances, although small, are reproducible across species (Fig. 2a). The most notable differences arise between reference-based and *de novo* OGC. Among the latter, the strategy for paralog discrimination determines the resulting OGC to a greater extent than the particular tools and identity thresholds, at least for the relatively permissive thresholds implemented in the study.

**Figure 2:**
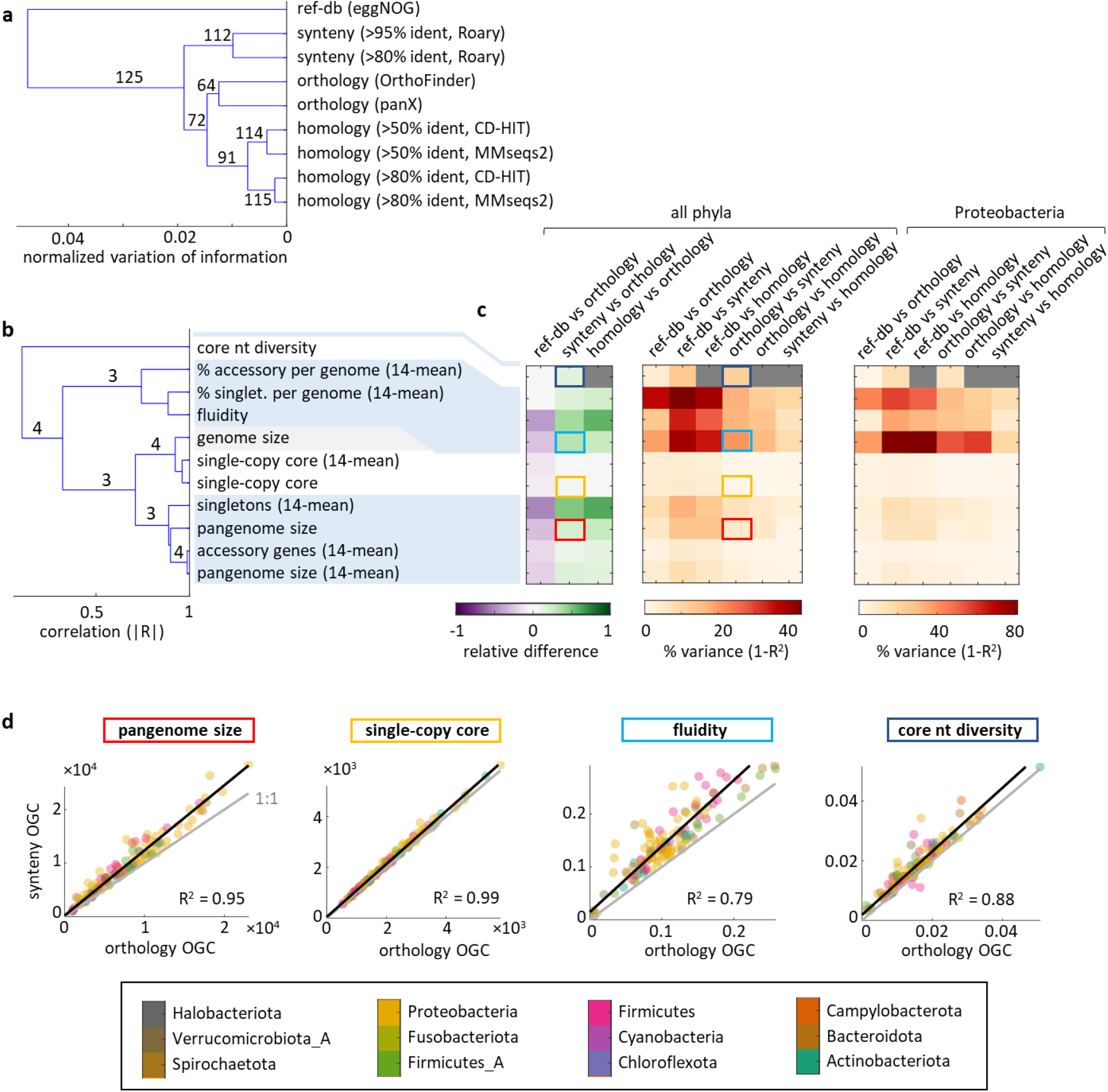
Method-dependent variation and uncertainty in pangenome features. (a) Consensus similarity tree of OGC building methods based on the species-wise normalized variation of information for the assignation of ORF to OGC. Labels indicate the number of species (out of 125) that support each branch. (b) Consensus similarity tree of different pangenome features based on pairwise, unsigned correlations. Labels indicate the number of OGC building methods (out of 4) that support each branch (values <3 are not shown). In all cases when the support is not complete, reference database mapping is the method that disagrees. (c) Quantitative comparison of pangenome estimates among methods. Left: relative differences in pangenome estimates; right: relative contribution of methodological choices to between-species variance. Note the different color scale for Proteobacteria. Highlighted cells correspond to the features shown in (d). (d) Species-wise comparison of selected pangenome features (pangenome size, number of single-copy core OGC, fluidity, and nucleotide sequence diversity in core genes) inferred from orthology- and synteny-based OGC. Black lines show the orthogonal least squares fit; gray lines indicate the 1:1 trend. Each point corresponds to the pangenome of one species, colored according to its phylum.

Based on the previous finding, we restricted our analysis to a single identity threshold (80%) and one tool of each class. We chose MMseqs2, panX, and Roary because they were primarily designed to study pangenomes or are at the core of popular workflows for pangenome analysis. Then, we investigated how paralog discrimination affects the properties of the inferred prokaryotic pangenomes. To that end, we selected 10 non-redundant quantitative features that represent different aspects of pangenome size and diversity. All *de novo* methods supported clustering these features in 4 groups (Fig. 2b) that can be interpreted in terms of (i) pangenome size, (ii) genome (and core genome) size, (iii) gene content diversity, and (iv) nucleotide diversity (evaluated in the core genes). As expected, reference-based OGC systematically produced the smallest estimates of pangenome size and gene content diversity (around 30% lower than orthology-based OGC), whereas *de novo* synteny- and homology-based OGC produced the largest estimates for those traits (20-35% higher than orthology-based OGC, see Fig. 2c).

Although the differences in some estimates (especially those involving singletons) are substantial, their practical impact on comparative analyses does not so much depend on their median magnitude, but on the extent to which they generate unexplained variation across species. More precisely, method-dependent variation appears if the choice of a particular set of OGC over another does not affect all species by the same factor. Because there is no ground truth for the “correct” set of OGC, method-dependent between-species variation constitutes an unavoidable source of uncertainty in comparative pangenome analyses that only becomes visible when considering multiple OGC construction methods. We assessed such uncertainty for 10 selected pangenome features by pairing sets of OGC (corresponding to different strategies for paralog discrimination) and quantifying the fraction of the total variance obtained with one set that remains unexplained after controlling for the values obtained with the other set. This measure also serves as a proxy for the methodological inconsistency in the estimation of pangenome properties. As shown in Fig. 2c-d (see also Suppl. Fig S1 and S2), core genome and pangenome sizes are generally consistent across methods; that is, using one method or another affects all species by the same factor. In consequence, methodological choices do not affect relative comparisons of core genome and pangenome sizes across species. In contrast, estimates of gene content diversity display higher levels of inconsistency, with method-dependent uncertainty accounting for roughly 20% of the total between-species variance (and up to 40% if using reference-based OGC). The inconsistencies in genome fluidity are especially high in the case of Proteobacteria, reaching around 50% of the total between-species variance.

Despite the known fact that sample size affects the estimation of pangenome properties, two lines of evidence prove that cross-method inconsistencies are independent of the number of genomes used to build the pangenome, at least within the range of 15-100 genomes explored in this study. First, the same trends are detected in gross and size-corrected features (for example, the total pangenome size and its average over subsets of 14 genomes). Second, cross-method scatter plots, like those of Fig. 2d, show no correlation between the number of genomes per species and the relative deviation of each species with respect to the regression line (Pearson’s correlation, |*R*| ≤ 0.16 and *q* > 0.05 for all features and OGC criteria included in Fig. 2c). Such lack of correlation indicates that method-dependent variation is equally likely to affect species with many or just a few available genomes.

A major limitation of reference-based approaches to OGC construction is that they are constrained by the limited diversity of the reference database. Indeed, in most species, 5-10% of the ORF could not be mapped to the eggNOG database and therefore were not assigned to any reference-based OGC (Suppl Fig. S3). The fraction of missing ORF is larger in some taxa that are underrepresented or absent from the reference database. The most extreme case, with >30% unmapped genes, corresponds to *B. burgdorferi*, the causal agent of Lyme’s disease. The poor performance of reference database mapping in *B. burgdorferi* is explained by the unique structure of its genome, which consists of a linear chromosome and >20 linear and circular plasmids without homologs in other species [39,40]. Despite these limitations, reference-based OGC provide reasonably good estimates for the number of core genes per genome and allow retrieving 85-90% of the single-copy core gene families identified by *de novo* approaches.

### Within-species paralogy and inference of core genomes

Among the pangenome features considered in this work, the size of the core genome is the least affected by the gene clustering criterion. However, a closer inspection of core gene families (defined in a strict sense, given the high completeness of the genomes included in the study) reveals differences in the mean copy number across methods (Fig. 3a). The fraction of core gene families that appear as single-copy ranges from 80% in reference-based to >99% in synteny-based OGC. In the case of reference-based and *de novo* homology-based OGC, the distribution of the mean copy number per core gene family per genome displays clear peaks at integer values, indicating the existence of complete sets of duplicated paralogs that were not resolved by these methods.

**Figure 3:**
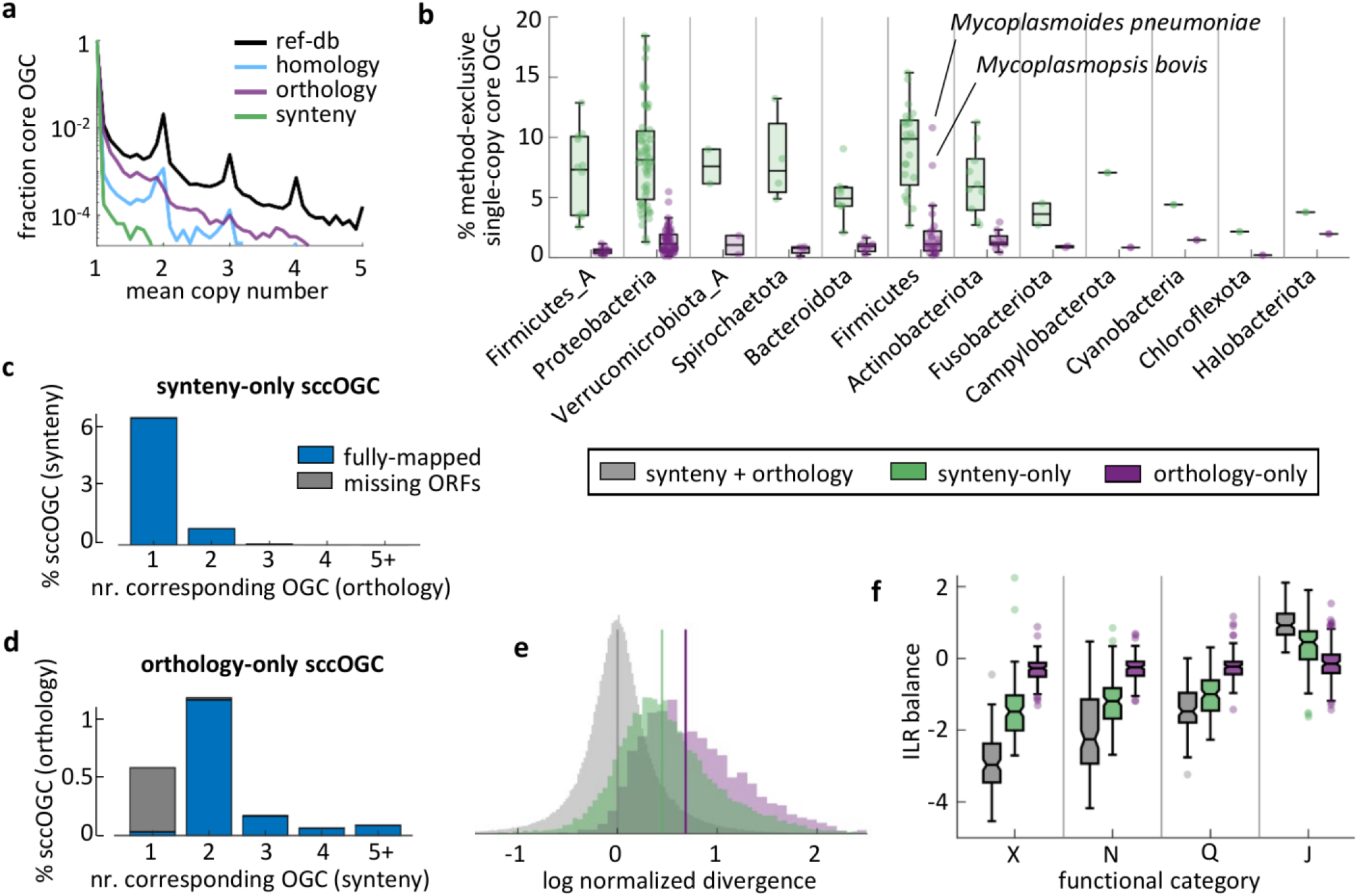
Comparison of core gene sets obtained from different OGC building methods. (a) Distribution of the mean copy number per core OGC per genome. (b) Fraction of all single-copy core (scc) gene families that are exclusively recovered through orthology- or synteny-based paralog splitting. (c) Distribution of synteny-exclusive, single-copy core OGC among orthology-based OGC. The large bar at “1” implies that most synteny-exclusive single-copy core OGC are subsets of larger orthology-based OGC that are core but not single-copy. (d) Distribution of orthology-exclusive, single-core OGC among synteny-based OGC. Most of the single-copy core OGC exclusively supported by orthology combine 2 accessory synteny-based OGC. In (c) and (d), gray bars denote method-exclusive single-copy core OGC that include sequences that could not be processed by the other method. (e) Nucleotide sequence divergence in single-copy core OGC exclusively supported by orthology, synteny, or both criteria. To account for between-species differences in evolutionary rates and phylogenetic tree spans, values were normalized by the species-level mean divergence (calculated from all single-copy core OGC that were supported by both synteny and orthology). Vertical lines indicate the distribution means for each group of single-copy core OGC. (f) Functional differences among single-copy core OGC supported by different criteria. Each set of box plots represents the balance (measured as the isometric log-ratio) between the relative frequencies of a given functional category (x-axis) and the remaining categories not considered in previous sets (e.g., the second set of box plots corresponds to the balance between functional category N and all the rest except X). The figure shows the 4 ILR balances with the greatest variation across methods. Abbreviations of functional categories, X: mobilome; N: cell motility; Q: 2° metabolites biosynthesis, transport and catabolism; J: translation, ribosomal structure and biogenesis. In (b) and (f), each data point corresponds to one species; boxes span the 25-75 percentiles; the central line indicates the median; whiskers extend to the most extreme data points that are not outliers; isolated points denote outliers; notches (only in f) show the 95% confidence interval of the median.

We next focused on single-copy core (s.c.c.) genes, which are of greater practical interest as they are often used to infer high-resolution phylogenies. Synteny-based methods systematically produce 5-10% more s.c.c. OGC than orthology-based methods (Fig. 3b). Typically, synteny-supported s.c.c. genes belong to orthologous OGC that are core but not necessarily single-copy (Fig 3c). Therefore, it appears that synteny criteria are effective in resolving single-copy representatives of core gene families affected by within-species duplications or HGT. On the other side, s.c.c. OGC that are orthologous but not supported by synteny tend to either contain incomplete ORFs (that could not be processed by the synteny-based workflow) or split among 2 or more synteny-supported accessory OGC (Fig. 3d). Such lack of synteny conservation could be a sign of non-vertical transmission in s.c.c. genes exclusively detected through orthology criteria. However, given the small fraction of these OGC in most species (<1% of all orthology-based s.c.c. OGC), the overall effect of such potential non-vertical contamination in downstream analyses is possibly modest. The only exception occurs in Mycoplasmatales, in which the relative contribution of non-synteny-supported OGC is amplified by their small genome sizes.

Single-copy core genes supported by a single criterion (either synteny or orthology) are significantly less conserved in terms of their average identity than those detected by both criteria (Fig. 3e); Kruskal-Wallis omnibus test *p* < 10^-20^). Among the s.c.c. genes that are method-exclusive, those based on orthology are significantly less conserved than those based on synteny (*p* < 10^-8^ for all Mann-Whitney post-hoc tests with Tuckey’s HSD correction). There are also significant differences in the functional profiles of s.c.c. genes that are supported by only one or both criteria (Fig. 3f), with overrepresentation of genes associated with mobile genetic elements, cell motility, and secondary metabolism, and underrepresentation of genes involved in translation among method-exclusive single-copy core genes (linear mixed effects model for isometric log-ratios; *F*(2,248) > 190, *q* < 10^-20^ in all cases).

### Systematic and species-specific biases in functional profiles

Estimates of genome content diversity are strongly affected by the choice of orthology- or synteny-based strategies to discriminate paralogs when building *de novo* OGC. To better understand the causes and implications of these differences, we classified ORF and OGC into 21 coarse-grained functional categories that are representative of the main molecular and cellular processes that take place in prokaryotic cells. For each functional category, we studied the agreement between orthology- and synteny-based OGC by calculating the NVI, the fraction of fully equivalent OGC (that is, OGC that contain exactly the same ORF regardless of the strategy for paralog discrimination), and the fraction of ORF assigned to fully equivalent OGC (Fig. 4a and Suppl. Fig. S4). By far, the highest inconsistency occurs for mobile genetic elements, for which only 25% of the OGC (encompassing 25% of the ORF) are equivalent. Besides mobile genetic elements, moderate degrees of inconsistency are observed for defense systems, intracellular trafficking/secretion, and replication/recombination/repair. On the other end, central cellular functions, such as translation, transcription, nucleotide metabolism, and coenzyme metabolism show the highest agreement, with 64-70% of the OGC (encompassing 77-84% of the ORF) being fully equivalent. These trends are also manifested when looking at the absolute and relative numbers of OGC per category (Fig. 4b), with synteny-based paralog discrimination producing a disproportional excess of OGC associated with the mobilome. The fraction of OGC that contain ORF from more than one functional category is consistently larger in the case of orthology-based OGC, although the absolute differences are modest (around 0.5-1% in most categories; Suppl. Fig S4). Functionally heterogeneous OGC are most often associated with signal transduction, cell cycle control/cell division, mobile genetic elements, and unknown or poorly characterized functions.

**Figure 4:**
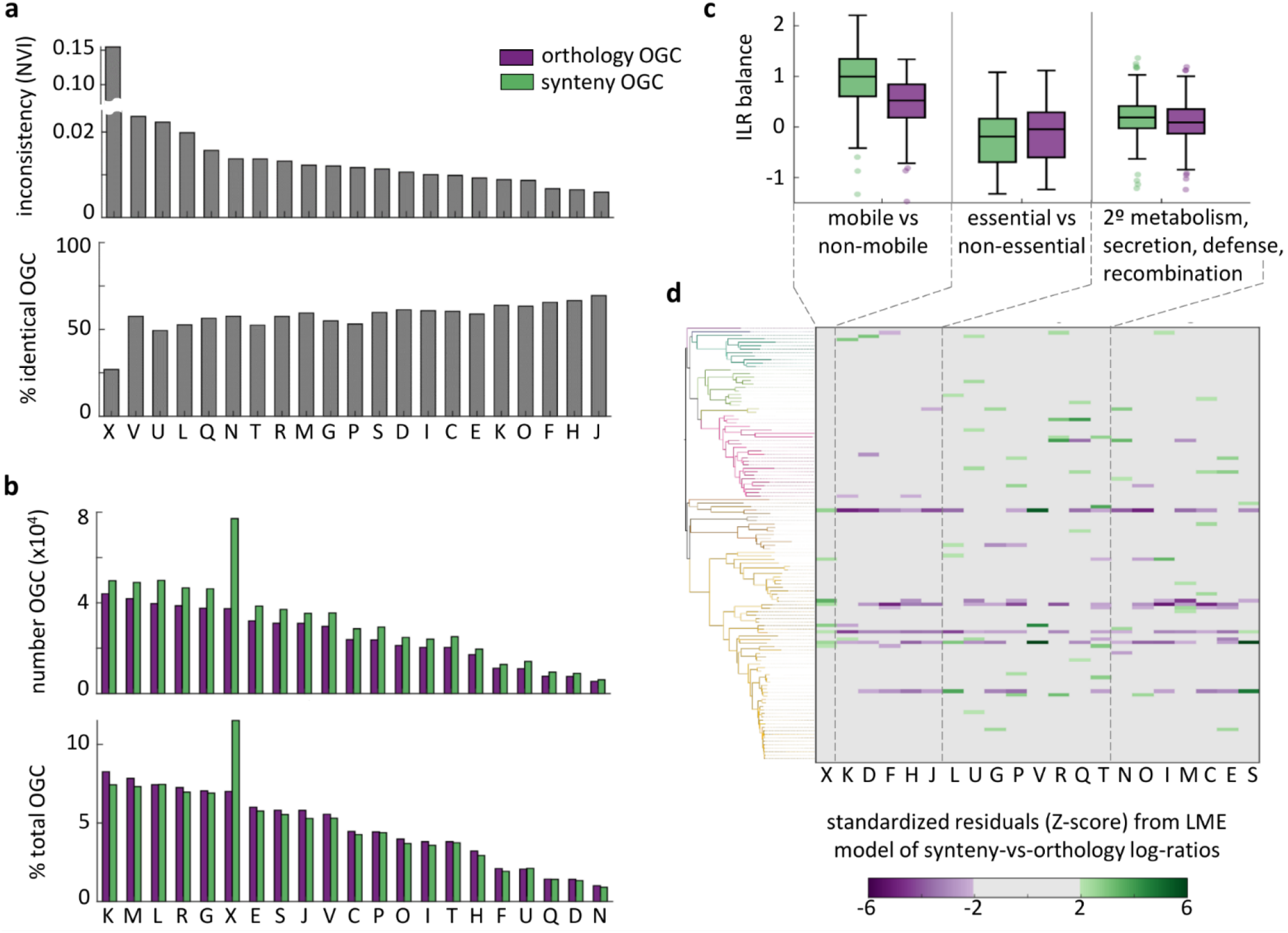
Systematic and specific biases in functional profiles associated with paralog splitting criteria. (a) Inconsistency of ORF assignations into OGC (normalized variation of information, top), and fraction of OGC that exactly contain the same ORFs (bottom) under synteny and orthology splitting criteria, stratified by functional category. (b) Absolute number (top) and relative fraction (bottom) of synteny- and orthology-based OGC associated with each functional category. (c) Balances (quantified as isometric log-ratios) for the functional categories that show the greatest systematic variation between paralog splitting criteria. Each set of boxplots represents the balance between the relative abundances of a group of functional categories (shown below) and all the remaining categories not considered in previous sets. Each data point corresponds to the pangenome of one species; boxes span the 25-75 percentiles; the central line indicates the median; whiskers extend to the most extreme data points that are not outliers; isolated points denote outliers. (d) Standardized residuals (Z-scores) of the linear mixed effects model used to infer the systematic differences shown in (c). Each row corresponds to the pangenome of one species, sorted according to the GDTB species tree [75] (phyla colored as in Fig 1). Colored cells indicate a significant excess of synteny-(green) or orthology-based (purple) OGC from a given category in a specific pangenome that is not explained by the general trends in (c). Abbreviations of functional categories, C: energy production and conversion; D: cell cycle control, cell division, chromosome partitioning; E: amino acid transport and metabolism; F: nucleotide transport and metabolism; G: carbohydrate transport and metabolism; H: coenzyme transport and metabolism; I: lipid transport and metabolism; J: translation, ribosomal structure and biogenesis; K: transcription; L: replication, recombination and repair; M: cell wall/membrane/envelope biogenesis; N: cell motility; O: posttranslational modification, protein turnover, chaperones; P: inorganic ion transport and metabolism; Q: secondary metabolites biosynthesis, transport and catabolism; R: general function prediction only; S: function unknown; T: signal transduction mechanisms; U: intracellular trafficking, secretion, and vesicular transport; V: defense mechanisms; X: mobilome: prophages, transposons.

Compositional analysis of functional profiles controlling for between-species variability confirms that synteny-based paralog discrimination leads to a significant increase in the fraction of OGC associated with mobile genetic elements (Fig. 4c; linear mixed effects model for isometric log-ratios; ILR-balance difference = 0.49, *F*(2,124) = 189, *q* < 10^-20^). Functional profiles derived from synteny and orthology-based OGC also differ in the balance between central cellular functions (transcription, translation, cell cycle, nucleotide and coenzyme metabolism) and other functional categories (ILR-balance difference = −0.11, *F*(2,124) = 132, *q* < 10^-20^); and between a set of functions including secondary metabolism, carbohydrate metabolism, secretion, defense and recombination, and the remaining functional categories (ILR-balance difference = 0.08, *F*(2,124) = 62, *q* < 10^-10^). Apart from these general trends, other significant differences between synteny- and orthology-based functional profiles are restricted to specific categories in one or a few particular species (Fig. 4d), such as defense in *Legionella pneumophila* (Z = 5.9, *q* < 10^-5^), *Borreliella burgdorferi* (Z = 5.4, *q* = 5×10^-5^) and *Bordetella pertussis* (Z = 4.4, *q* = 0.002), secondary metabolism in *Bacillus anthracis* (Z = 4.0, *q* = 0.008), and signal transduction in *Brachyspira hyodysenteriae* (Z = 3.8, *q* = 0.019).

### Variability of gene flux estimates

To assess whether method-dependent variation in pangenome composition propagates to downstream analyses, we investigated the effect of synteny- and orthology-based paralog discrimination on a quantitative study of genome dynamics. To that purpose, we used the software Gloome, that infers events of gene gain and loss along a lineage taking as inputs the strain-level phylogenetic tree and a binary matrix with the presence and absence profiles of each OGC. As shown in Fig. 5a, using synteny-based instead of orthology-based OGC leads to a 60% increase in the estimated number of gene gains and losses per lineage. Genome vs gene change ratios, measured as the number of expected gains and losses per gene per core nucleotide substitution are also higher (40% increase) when using synteny-based OGC. In contrast, the ratio between gene gains and losses, that determines the short-term dynamics of genome size [41], displays a more complex response, with synteny-based OGC producing lower or higher estimates than those obtained with orthology-based OGC depending on whether a species is dominated by gains or losses. Method-dependent uncertainties account for 15%, 18%, and 30% of the between-species variability in the total flux, the genome vs gene change ratio, and the gain vs loss ratio, respectively. These results indicate that comparative analyses of short-term genome dynamics are highly sensitive to methodological choices for paralog discrimination, especially when it comes to evaluate the balance between gene gain and loss.

**Figure 5:**
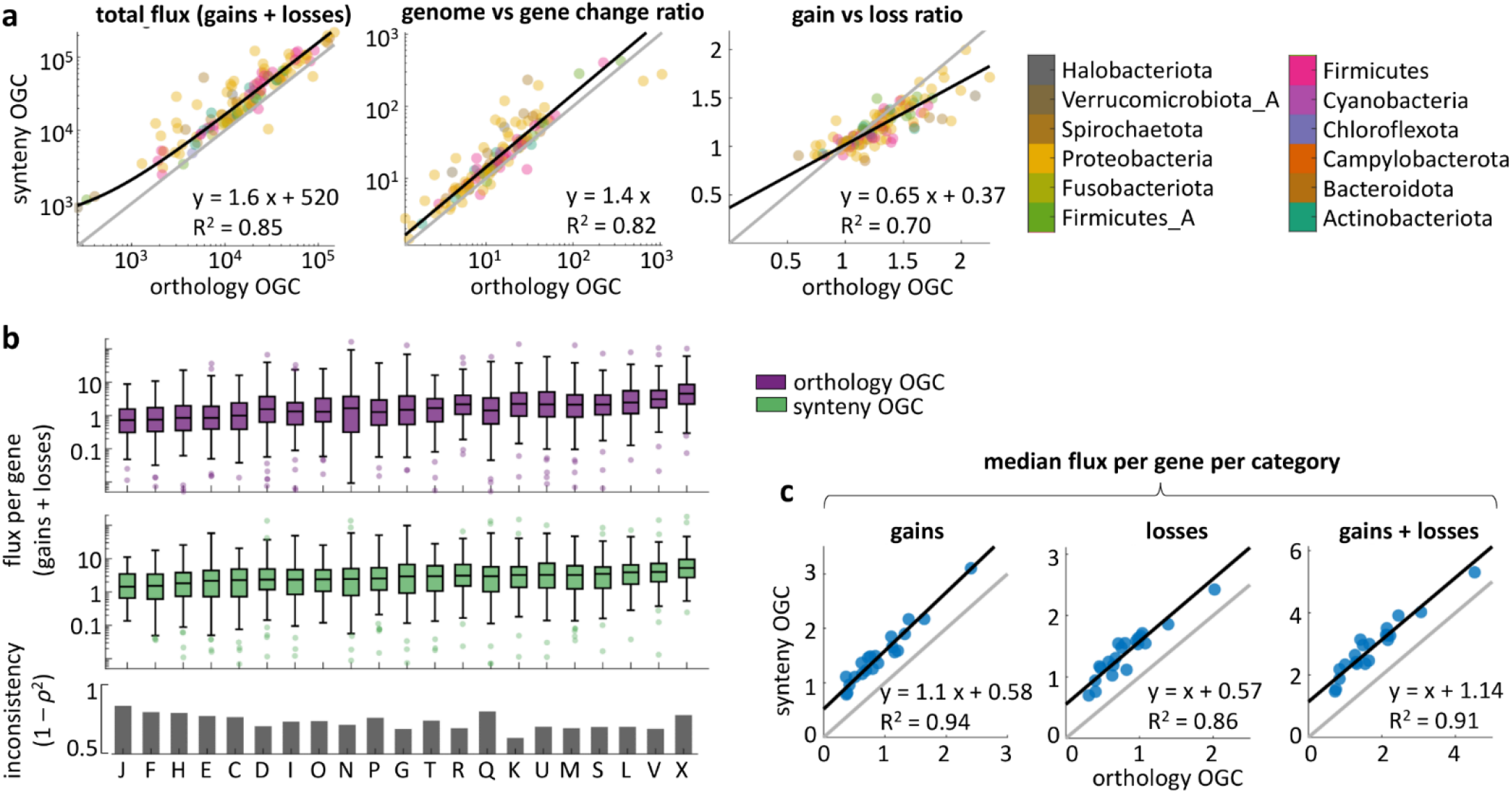
Effect of paralog splitting criteria on the inference of gene flux. (a) Species-wise comparison of the total gene flux (gains and losses along the species tree, left), genomic content vs gene sequence evolution ratio (middle), and total gain vs loss ratio (right) inferred from synteny- and orthology-based OGC. The method-dependent uncertainty is equal to 1 – R^2^. (b) Flux per gene per functional category inferred from orthology-(top) and synteny-based (middle) OGC. Each data point corresponds to one species; boxes span the 25-75 percentiles; the central line indicates the median; whiskers extend to the most extreme data points that are not outliers; isolated points denote outliers. The bar plot at the bottom shows the inconsistency between methods, quantified as one minus the squared rank correlation. Abbreviations of functional categories as in Fig. 2. (c) Median flux per gene per category, calculated over all the species, for orthology-based (x-axis) and synteny-based (y-axis) OGC. Similar trends are observed for gains, losses, and the combination of both.

A deeper analysis of gene flux by functional categories reveals that inconsistencies between gene clustering criteria (quantified as one minus the squared rank correlation of species-wise estimates to account for the numerous outliers) are stronger among mobile genetic elements and genes involved in secondary metabolism and inorganic ion transport (Fig. 4b). Although high inconsistencies are also observed in central functional categories, such as translation, the practical relevance of those is lesser due to the relatively low fluxes associated with those categories. If all species are jointly considered by calculating their median, flux estimates obtained from synteny OGC display a systematic deviation of >1 additional event per gene in all functional categories, which is evenly distributed between gains and losses (Fig. 4c).

### Estimation of pangenome properties from medium- and low-quality genome assemblies

Pangenomes built from high-quality, nearly complete genomes represent an appropriate benchmark to compare gene clustering criteria without the confounding effects of genome incompleteness and contamination. However, complete genomes are relatively rare and unevenly distributed among taxonomic groups. Given those limitations, incomplete genomes, often assembled from metagenomes, constitute the only available choice to study the pangenomes of many bacteria and archaea. To evaluate the performance of different gene clustering criteria in such suboptimal scenarios, we built 96 alternative pangenomes by replacing some of the complete genomes by an increasing proportion of medium- and low-quality genomes assembled from metagenomes (MAG). To keep the computational cost manageable, we focused on 4 species belonging to the phyla Proteobacteria, Actinobacteria, Firmicutes, and Bacteroidota, that cover a broad range of pangenome sizes and gene content diversity. For each pangenome feature, we quantified the uncertainty introduced by MAG as the relative amount of variation in the alternative pangenomes that could not be predicted from the original high-quality pangenomes. Values greater than 1 (that correspond to R^2^ < 0) imply that pangenomes that include MAG follow qualitatively different trends than those built from complete genomes. In those cases, the inclusion of MAG leads not only to loss of precision, but to systematic biases.

Estimates of pangenome size are relatively unaffected by the inclusion of medium- and low-quality MAG, especially if pangenome size is calculated by averaging over multiple subsamples (Fig. 6a, top; Suppl. Fig. S5a). By using reference, homology, or synteny OGC, the unexplained variance in pangenome size can be kept below 10% even if half of the genomes are incomplete. In contrast, estimates of gene content diversity are much more sensitive to genome incompleteness (Fig. 6a, middle; Suppl. Fig. S5a and S6), with systematic deviations in fluidity appearing with only a 5% of MAG. As expected, incomplete genomes severely impair the direct detection of core genes (Suppl. Fig. S7), although consistent estimates of the core genome size (with up to 50% medium- and low-quality MAG) can be obtained with mOTUpan [42], a specialized tool that uses Bayesian inference techniques to predict core genes while accounting for genome incompleteness (Fig 6a, bottom; Suppl. Fig. S5a).

**Figure 6:**
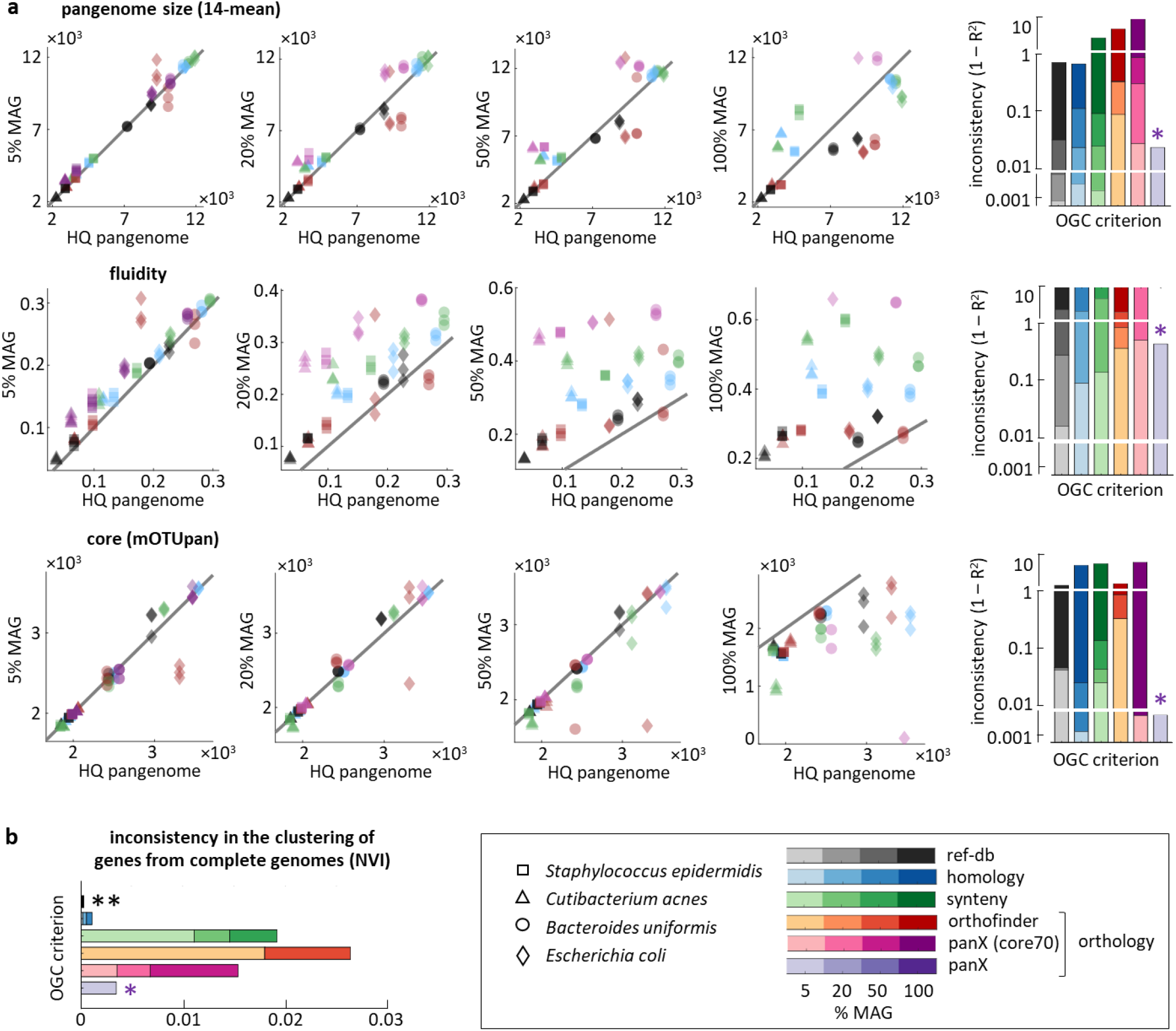
Estimation of pangenome properties from incomplete genomes. (a) Comparison between pangenome properties inferred from high-quality genomes (x-axis) and mixtures of medium-quality MAG and high-quality genomes (y-axis). From left to right, the scatter plots correspond to pangenomes with 5%, 20%, 50% and 100% of MAG. Each point in the scatter plots corresponds to one pangenome, with different symbols and colors used to distinguish among species and OGC generation methods, respectively. Each scatter plot combines data from 4 species, 6 gene clustering methods, and 3 random subsamples. The bar plots on the right summarize the observed inconsistencies, with color intensities representing the fraction of MAG. Note that panX fails to produce results in pangenomes that contain >5% of MAG (purple asterisks). (b) Sensitivity of gene clustering methods to the addition of MAGs, calculated by comparing the cluster assignations of genes from high-quality genomes before and after adding MAG. NVI: normalized variation of information. Different OGC generation methods are color coded, with color intensities indicating the fraction of MAG. Note that reference-database mapping methods produce an NVI equal to zero (double asterisk).

Inspired by the good performance of mOTUpan, we explored the potential of Bayesian approaches to produce more robust measures of gene content diversity. Thus, we recalculated the fraction of accessory genes per genome by combining mOTUpan’s predictions for the number of core genes and the mean genome size (see Methods). The new metrics is more consistent than fluidity, although still far from reaching the high robustness of pangenome size estimates (Suppl. Fig. S6). Taken together, the higher performance of mOTUpan for the estimation of core genes and this “mOTUpan-based” metrics for gene content diversity illustrate how a probabilistic framework can help overcoming some of the limitations imposed by incomplete and poorly assembled genomes.

To better understand variations in performance across methods, we looked at genes from complete genomes and studied whether the inclusion of MAG leads to their being reassigned to different clusters (Fig. 6b; Suppl. Fig. S5b). Because reference-based methods operate gene by gene, the addition of new genomes does not have any effect on how a given gene is clustered. Such “structural” robustness explains why reference-based OGC provide the most consistent estimates in the presence of MAG. In stark contrast, clustering based on synteny and orthology is substantially affected by the addition of incomplete or poorly assembled genomes (the within-method NVI after adding 5% of MAG becomes as large as the cross-method NVI in high-quality pangenomes, shown in Fig. 4a). The fact that this phenomenon is not observed for homology-based OGC indicates that the paralog discrimination algorithms used by synteny- and orthology-based methods are prone to misclassify genes from high-quality genomes if the dataset also includes lower-quality genomes.

## Discussion

Orthology is generally considered the optimal criterion for clustering gene sequences for comparative genomics [14]. Because orthologs tend to conserve their function and evolve consistently with speciation patterns, they are, at least in theory, the most fitting choice for functional and phylogenomic studies. In practice, however, the use of orthology as a gold standard for pangenome analysis faces technical and conceptual challenges. On the technical side, distinguishing orthologs from paralogs is computationally costly. Therefore, to deal with large (meta)genomic datasets, orthology prediction tools often resort to heuristic algorithms that risk missing true orthologs or including out-paralogs (paralogs that duplicated before the last common ancestor of the clade of interest). On the conceptual side, intra-species gene duplication and horizontal gene transfer generate multi-copy gene lineages that, despite not violating the orthology criterion at the species level, may confound downstream analyses that assume vertical transmission. In those cases, a stricter criterion that only clusters together vertically transmitted members of a gene family may be more desirable.

Motivated by these caveats, we assessed how three major clustering criteria (homology, orthology, and synteny) implemented by five popular pangenome analysis tools affect pangenome reconstructions and downstream phylogenomic analyses. Although we only tested a limited number of tools and parameter settings, our results suggest that the underlying formal criterion for paralog discrimination (shared ancestry at speciation for orthology, conserved gene neighborhood for synteny), rather than the actual implementation, is what drives qualitative differences across methods.

Previous works had pointed out that single-species estimates of pangenome size and diversity strongly depend on the method used to cluster genes [19–21]. We confirmed such observations and expanded on the implications for comparative pangenome analyses. When conducting comparative studies, inconsistencies in relative differences and cross-species trends are of much greater concern than absolute differences in single-species estimates. In that regard, trends involving pangenome and core genome sizes are generally robust across methods. More precisely, choosing one or another method affects all species-wise size estimates by the same constant multiplicative factor. In contrast, cross-species comparisons of genome plasticity and pangenome diversity are highly sensitive with respect to the clustering criterion, in a way that cannot be explained by any linear or nonlinear data transformation. Such inconsistencies in pangenome diversity, that reach up to 50% of the total between-species variance in Proteobacteria, should be a major source of concern for studies aimed at understanding the forces that shape microbial pangenomes. For comparison, it has been estimated that habitat and phylogeny contribute to explain approximately 20% of the between-species variance in pangenome diversity [6]. That said, regardless of the method used to discriminate paralogs, the lowest rates of gain and loss are always observed in genes associated with central cell functions, whereas the highest rates correspond to mobile genetic elements and defense systems. Therefore, the inverse association between gene flux and essentiality described by previous studies [43–46] appears as a robust feature of genome plasticity.

The contribution of a particular gene clustering method to pangenome variability can only be assessed by comparison with other methods, which is often unfeasible in large datasets. As a result, methodological “noise” remains unnoticed, potentially contributing a relevant fraction of the unexplained variance or, in the worst-case scenario, acting as a confounding factor if methodological biases correlate with the biological variables of interest.

Technical choices during genome assembly and ORF prediction can also cause biases that propagate downstream affecting pangenome analyses [35]. For example, variations in assembly completeness and contamination will have a direct repercussion on the fraction of core genes and genes with no homologs in the pangenome. By studying the interplay between assembly quality and gene clustering criteria, we found that reference-based clustering produces the most robust estimates of pangenome properties in pangenomes that include incomplete and poorly assembled genomes. On the other side, orthology-based methods often lead to inconsistent results even with relatively low proportions of medium-quality assemblies. The most likely cause for such poor performance is that incomplete genomes interfere with the construction of the strain-level reference trees that serve as guides for the identification of orthologs. For example, the default for panX is to build guide trees from strict core genes, so that the method fails if no core genes are found (which typically occurs if >5% of all genomes are incomplete). More permissive methods that take into account the “soft” core, such as OrthoFinder or panX with customized core thresholds, can handle higher proportions of incomplete genomes. However, their accuracy will still be compromised if the guide tree is incorrect.

### Practical guidelines

Our findings stress that, as long as choosing an operational definition of gene cluster remains arbitrary, pangenome properties affected by method-dependent variability will be subject to intrinsic uncertainty. To minimize such uncertainty, the selection of tools for gene clustering should be primarily guided by the nature of the research goals, rather than by computational considerations such as runtime and memory usage. In the next paragraphs, we provide some practical guidelines to help selecting optimal clustering criteria for different purposes. A summary of those guidelines can be found in Table 2.

**Table 2:** Practical guidelines to select optimal gene clustering criteria for pangenome analysis.

| OGC | Strengths | Limitations | Recommended use |
| --- | --- | --- | --- |
| Ref-db | - Annotation and classification at once.<br>- The most robust method in pangenomes with incomplete genomes. | - Loss of all gene families that are not represented in the ref-db.<br>- Discrimination of paralogs is limited by the taxonomic resolution of the ref-db. | - Functional profiling and detection of single-copy core genes (if missing taxon-specific genes is not a problem).<br>- Analysis of pangenomes with a large fraction of incomplete genomes.<br>- Cross-species tracking of pangenome contents. |
| Homology | - Faster than any other method. | - Setting a hard similarity threshold can lead to merging paralogs and/or missing distant orthologs. | - Quick detection of single-copy core genes.<br>- Fast de novo clustering of sequences without homologs in ref-db.<br>- Analysis of very large pangenomes. |
| Orthology | - Evolutionary consistent clustering of mobile gene families.<br>- Discrimination of in- and out-paralogs. | - Computationally costly (panX becomes prohibitive with >200 genomes).<br>- Poor performance if the pangenome contains incomplete genomes. | - Unbiased functional profiling.<br>- Study of genome plasticity and gene flux in high-quality pangenomes.<br>- Assessment of within-species gene expansions. |
| Synteny | - Accurate discrimination of vertically transmitted gene families.<br>- The most sensitive method for single-copy core genes. | - The fragmentation of mobile gene families in multiple OGC can bias functional profiles and quantitative estimates of genome plasticity. | - Identification of vertically transmitted orthologs.<br>- Generation of high-resolution phylogenetic trees.<br>- Analysis of pangenomes with up to 20% of incomplete genomes. |

The identification of single-copy core genes is one of the main outcomes of pangenome reconstruction [47]. Single-copy core genes allow building high-resolution strain trees that can be very robust to the effects of unbiased homologous recombination [48]. However, to obtain accurate trees, it is fundamental that the OGC that correspond to single-copy core genes are not contaminated by HGT [10]. As expected, synteny is the most sensitive criterion to discriminate single-copy representatives among gene families affected by paralogy and intra-species duplications. Nevertheless, *de novo* homology- and orthology-based methods retrieve most (90-95%) synteny-supported single-copy core OGC with low (<1%) potential contamination, which makes them good alternatives for many practical applications. Regardless of the criterion used to obtain the OGC, if the pangenome includes incomplete genomes, inference of core genes can be notably improved by applying probabilistic approaches, such as those implemented by mOTUpan [42].

Synteny-based gene clusters display substantial fragmentation of non-core gene families, that affects not only mobile genetic elements but also genes involved in defense, secondary metabolism, signal transduction, and secretion, among others. Because such fragmentation is not homogeneous across taxa and functional categories, synteny-based clustering can introduce biases in comparative studies of pangenome function and dynamics. Functional profiles based on orthology (rather than synteny) are better choices, as they are more robust with respect to the amplification of mobile genetic elements and other accessory gene families that are characteristic of some species [22,49,50]. Likewise, inference of gene fluxes within and across pangenomes with tools that account for gene copy number (e.g., the phylogenomic software COUNT [27]) should be performed with orthology-based gene clusters. In turn, analyses of genome plasticity based on binary (presence or absence) phyletic profiles may be more accurate if using synteny-based gene clusters, because they better capture the contribution of intra-species paralogs to the total gene flux. Future studies aimed at exploring the role of adaptive and non-adaptive processes in pangenome evolution should explicitly discuss how methodological choices affect the interpretation of their results. For example, choosing synteny over classical orthology as the clustering criterion may inflate the proportion of mobile genetic elements in the pangenome, altering the balance between selfish and potentially beneficial accessory genes. Estimates of gene content diversity are extremely sensitive to the presence of incomplete genomes, even if they only constitute a small fraction of the whole pangenome. Therefore, to study pangenome plasticity, it is preferable to work with a reduced set of high-quality genomes than with a larger set of lower-quality genomes. If incomplete genomes cannot be avoided, we recommend using reference-based OGC and measuring gene content diversity as the predicted (rather than observed) percentage of accessory genes per genome.

As a fast alternative to *de novo* clustering methods, we evaluated the performance of OGC built by annotating gene sequences with the eggNOG reference database. Although reference-based OGC deviate from *de novo* OGC in many aspects, they are a reasonable option to identify single-copy core genes and generate functional profiles in well-studied genera, especially if paralogs and rare genes can be disregarded. Reference-based OGC are also the most robust option to deal with pangenomes that contain a large fraction of incomplete or poorly assembled genomes. On the other side, eggNOG-based OGC can miss a significant portion of species-specific core genes in poorly sampled taxa, do not discriminate paralogs below the genus level, and filter out gene families with narrow taxonomic distribution. Accordingly, pangenome analyses using reference-based OGC should justify their conclusions accounting for these limitations. Possibly, the most interesting application of reference-based OGC is found in large-scale comparative studies that require tracking OGC across species. That can be achieved thanks to the hierarchical structure of the eggNOG database, that allows annotating orthologs at multiple taxonomic levels without additional computational cost.

## Conclusions

There is no “one size fits all” gene clustering method for pangenome analyses. Such lack of a universal method does not only reflect the practical limitations of current algorithms, but a deeper conceptual trade-off. For applications that involve tracking vertically transmitted genes, the best OGC are those that effectively discriminate among in-paralogs, which is generally achieved by applying synteny criteria. However, those same OGC are not optimal to study within-species expansions and contractions of accessory gene families. (For that purpose, OGC should keep in-paralogs together, as dictated by classical orthology.) In practice, the best choice also depends on the quality of the genomic assemblies, with homology and reference-based OGC producing the most robust results in medium- and low-quality pangenomes.

Reusability and meta-analysis of pangenome datasets is currently hindered by the incompatibility of OGC obtained by different methods. To address that limitation and foster future research, it is critical to come to a consensus on a set of methods that cover the most relevant criteria for paralog discrimination. To maximize consistency and reusability, an optimal solution should leverage the hierarchical nature of gene clustering criteria, with synteny OGC nested within orthology OGC, and the latter nested within homology OGC. Such multi-level scheme would facilitate selecting the best set of OGC for each purpose and assessing the robustness of pangenome properties in the face of methodological uncertainty. We expect that the multi-method dataset generated in this study (publicly available in https://dx.doi.org/10.5281/zenodo.7387758), though not strictly nested, will constitute a first step in that direction.

## Methods

### High-quality genomic sequences

We parsed the metadata files of the Genome Taxonomy Database (GTDB) release 95 [51] to identify all the species (sensu GTDB) for which there were at least 15 high-quality available genomes. High-quality genomes were defined according to the MIMAG criteria [52] plus the following more stringent filters: completeness >99% and contamination <1%, both assessed through CheckM [53], mean contig length >5kb, and contig count <500. Metagenome-assembled genomes (MAG) and single-amplified genomes (SAG) were not included in the analysis. After applying these filters, we recovered 321 bacterial and 1 archaeal species, belonging to 125 different genera (sensu GTDB). To minimize possible taxonomical biases, only one representative species per genus was selected for subsequent analyses. For those genera with >1 suitable species, we kept the species with the highest number of high-quality genomes, as they presumably were the most informative for pangenome studies.

To keep our analyses within a manageable computational cost, species with >100 high-quality genomes were subsampled to keep at most 100 genomes per species. To that end, we separately aligned the amino acid sequences of 120 nearly universal marker genes employed by the GTDB with MAFFT [54], concatenated the alignments, and ran IQ-Tree [55] to obtain phylogenetic trees including all the strains of the same species. Then, we applied a heuristic subsampling strategy that maximized the diversity of the subsampled genomes whilst accurately reflecting the topology of the strain trees.

The final dataset comprised 6,796 bacterial genomes belonging to 124 species and 55 archaeal genomes belonging to 1 species. The genomes were downloaded from the NCBI FTP site. Open reading frames (ORF) were predicted with Prodigal v2.6.3 [56], using the “single” mode that is recommended for finished genomes and quality draft genomes. All runs used the bacterial, archaeal and plant plastid code 11 (https://www.ncbi.nlm.nih.gov/Taxonomy/Utils/wprintgc.cgi), which maps UAA, UGA, and UAG to stop codons, except for *Mycoplasmopsis bovis PG45* (GCF_000183385.1) and *Mycoplasma pneumoniae FH* (GCF_001272835.1), which used code 4 and translate UGA to tryptophan.

### Metagenome-assembled genomes

Medium- and low-quality MAG for *Escherichia coli*, *Cutibacterium acnes*, *Bacteroides uniformis*, and *Staphylococcus epidermidis* were downloaded from the Global Microbial Genomic Bins database (GMBC 1.0) [57], and their ORF were predicted with Prodigal using the same options as with high-quality genomes.

For each species, we built 24 mixed pangenomes organized in two series, one for low-quality MAG and the other for medium-quality MAG. Each series consists of 12 pangenomes, with 5%, 20%, 50%, and 100% MAG (the rest being high-quality genomes), and three random replicates for each composition. To build each pangenome, we randomly ampled the required number of genomes from the collection of MAG and high-quality genomes. All mixed pangenomes contained a total of 100 genomes, except for *Bacteroides uniformis*, for which only 65 high-quality genomes were available, and *Staphylococcus epidermidis,* for which only 88 and 84 medium and low quality MAGs were available, respectively.

### De novo OGC construction

We generated eight sets of *de novo* species-level OGC, each one representing an alternative approach to identify homologous genes and discriminate within-species paralogs (Table 1). To facilitate comparison among methods, we started in all cases from the same predicted ORF previously obtained with Prodigal, overriding any optional ORF prediction step provided by the OGC construction software.

Four sets of homology-based OGC were built with the sequence clustering tools MMseqs2 [29] and CD-HIT [28], setting the minimum identity threshold to 50% and 80%. MMseqs2 was run with the options ‘easy-cluster’ for cascaded clustering and ‘cluster-mode 1’ to define clusters based on connected components. The options for CD-HIT were set so that sequence homology was calculated locally, the alignments covered >80% of the longest sequence, and sequences were assigned to the best-matching cluster (-G 0 -aL 0.8 -g 1 -M 8000). The word size for CD-HIT (option -n) was set to 5 for minimum identity 80% and 3 for minimum identity 50%, as recommended by the developers. MMseqs2 and CD-HIT do not perform any kind of paralog splitting; therefore, the resolution of the resulting OGC only depends on the similarity threshold used for clustering.

Two sets of synteny-based OGC were built with the software Roary [33], setting the minimum identity threshold to 80% and 95%. The Roary algorithm starts by pre-clustering highly-similar protein sequences with CD-HIT [28] to obtain a smaller set of representative sequences. Roary then conducts an all-against-all comparison with BLAST and filters the hits based on the user-provided identity threshold. Based on the network of hits, representative sequences are clustered with MCL [58] and the resulting clusters are merged with the pre-clusters. As a final step, Roary uses conserved gene neighborhood information to split homologous groups containing paralogs into groups of synteny-supported OGC.

Two sets of orthology-based OGC were built with the software panX [32] and OrthoFinder [31]. Both algorithms initially cluster sequences in orthologous groups by performing an all-against-all similarity search with DIAMOND [59] and posterior clustering with MCL. The hits retrieved by DIAMOND are filtered only in terms of statistical significance, regardless of sequence identity, which allows recovering relatively divergent homologs. In the panX algorithm, the sequences of these initial clusters are aligned with MAFFT [54] and cluster-level phylogenetic trees are built with FastTree [60]. Finally, panX obtains orthology-supported OGC by examining the resulting trees and applying a set of heuristic rules to split paralogs from true species-level orthologs. OrthoFinder directly builds phylogenetic trees from the DIAMOND scores, infers the root based on gene duplication patterns, and generates OGC that are compatible with the rooted trees.

### OGC construction by reference database mapping

Reference-based OGC were built by mapping translated ORF to the eggNOG (evolutionary genealogy of genes: Non-supervised Orthologous Groups) database version 5.0 [61]. For that purpose, we ran eggNOG-mapper v2 [62] with command line options “*-m diamond --tax_scope_mode narrowest*” to search queries against eggNOG sequences with DIAMOND and transfer orthologous group annotations at the highest possible taxonomic resolution (which typically corresponds to the genus level).

### Pangenome features

OGC presence-absence matrices (sometimes known as phyletic profiles) and gene-to-OGC relationships were used to estimate a set of pangenome features that intend to capture both the size and the diversity of the pangenome.

Pangenome size was calculated as the total number of OGC retrieved for each species. Because this measure is positively correlated with the number of genomes sampled, an unbiased estimate was obtained by randomly subsampling sets of 14 genomes and taking the average over 100 realizations (we refer to this and other pangenome features obtained with the same procedure as “14-mean”). To ensure that the results obtained with these two measures were sufficiently representative, we calculated three additional measures of pangenome size: Chao’s lower bound [63]; the normalized pangenome size, obtained by dividing the pangenome size (*P*_*tot*_) by the sum of the harmonic series of the number of genomes (*n*), such that 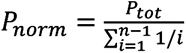 [64]; and the pangenome size of the 15 most dissimilar genomes in terms of genome content. After verifying that all those metrics were highly correlated across species (R>0.95), we proceeded with the uncorrected and 14-mean pangenome sizes, which are simpler and more easily interpretable.

We defined the core genome as the set of OGC contained in all the genomes of a species. This strict definition is appropriate given the almost full (>99%) completeness filter imposed on high-quality genomes. Pangenomes were also characterized in terms of the absolute number of single-copy core OGC, accessory OGC (those that are not core), and singleton OGC (those encompassing a single ORF). As measures of genome content diversity, we computed the mean percentage of OGC that are accessory and singleton with respect to all the OGC present in each genome, averaged over 100 random subsamples of 14 genomes per species. In addition, a measure of genomic fluidity that quantifies the average dissimilarity in gene content of randomly sampled pairs of genomes [65] was obtained with the ‘fluidity’ function of the R package micropan [66]. To ensure that variations in fluidity were not simply due to the randomness of the subsampling procedure, we conducted a series of preliminary tests to determine the minimum number of sampled pairs required for convergence in fluidity estimates. We found that using 500 random pairs (instead of only 10 pairs that is the default in micropan) reduced the relative variability across repeated estimates below 0.5%. Therefore, we set the option ‘n.sim = 500’ for all fluidity calculations.

The species-wise core gene alignment similarity (CGAS) was calculated using ORFs that belong to single-copy core OGC. Pairwise global alignments of all the ORFs assigned to the same OGC were performed with the Needleman-Wunsch algorithm as implemented by the *needleall* program of the EMBOSS suite [67]. After removing all self-alignments (alignments of ORFs with themselves), the average identity of each pair of genomes was calculated as ∑ *m*_*i*_ / ∑ *L*_*i*_ where *m*_*i*_ and *L*_*i*_ represent the number of matches and total alignment length for the pair of sequences from the i-th OGC and the sum extends over all single-copy core OGC. The species-wise CGAS was obtained as the average over all pairs of genomes of the same species. The nucleotide sequence divergence in single-copy core genes was defined as one minus the species-wise CGAS. Given the high computational cost of this procedure, calculation of CGAS and core gene divergences was restricted to reference-, synteny-, and orthology-based OGC.

### High-resolution species trees and inference of gene gains and losses

High resolution trees were obtained for each species using concatenated alignments of all single-copy core OGC obtained by Roary. Amino acid sequences for each single-copy core OGC were aligned with mafft-linsi (L-INS-I Algorithm, default options) [54] and back-translated to nucleotide alignments with the program pal2nal.pl using codon table 11 (except for species of the order Mycoplasmatales, which use codon table 4) [68]. The nucleotide alignments were concatenated and used as input for the tree construction program FastTree (command line options -gtr -nt -gamma - nosupport -mlacc 2 -slownni) [60]. The preliminary tree produced by FastTree was subsequently provided to RAxML for branch length optimization (options -f e -c 25 -m GTRGAMMA) [69]. The package ETE3 [70] was used for mid-point rooting and visualization.

Gene gains and losses along each species tree were inferred with the phylogenomic reconstruction software Gloome [71]. As the input for Gloome, we used the phyletic profiles for the presence or absence of each OGC and the high-resolution species trees; options were set to optimize the tree branch lengths under a genome evolution model with 4 categories of gamma-distributed gain and loss rates and stationary frequencies at the root. To compare among OGC generation methods, we varied the input phyletic profiles according to the desired method while keeping the species trees unchanged. The Gloome optimization algorithm did not converge for *Enterobacter himalayensis* and *Chlamydia muridarum*; therefore, those species were excluded from all the analyses involving gene gains and losses.

### Functional annotation and statistical analysis of functional profiles

Functional annotation at the gene level was done by mapping individual genes to custom-made HMM profiles of the 2020 release of the COG database [72]. Functional annotation at the OGC level was done by applying the majority rule to gene-level annotations. Coarse-grained pangenome functional profiles were built by counting the number of OGC assigned to each of the 21 major prokaryotic functional categories defined in the COG database.

The statistical analysis of functional profiles was conducted by applying the phylofactorization framework [73]. First, to account for the constant-sum constraint that complicates the statistical analysis of compositional data, we applied the isometric log-ratio (ILR) transform to the species-wise functional profiles. Informative ILR balances were defined following a guide tree that was obtained by calculating the mean differences between synteny and orthology OGC for each functional category and performing hierarchical clustering of the functional categories based on such differences. Then, linear mixed effects models were set out for each of the 20 ILR balances, with OGC clustering criteria as fixed effects and species as random effects. Model fitting was performed with the R package lmerTest [74] and contrasts among balances were conducted with the R package phylofactor (https://github.com/reptalex/phylofactor), which ranks the balances based on the fraction of the total variance that is explained by the model. Statistical significance was calculated by using Satterthwaite’s approach to estimate the degrees of freedom of the F-statistic and applying Bonferroni correction to account for multiple comparisons.

### Comparison of high-quality and MAG-derived pangenomes

To quantify the inconsistency between pangenomes built from high-quality genomes and pangenomes built from mixtures of MAG and high-quality genomes, we calculated the fraction of unexplained variance in the latter that cannot be explained by the former. If the value of a given pangenome feature in a given species is represented as a pair (*x*, *y*), where *x* and *y* represent, respectively, the value in high-quality and mixed pangenomes, then the fraction of unexplained variance is equal to ∑(*y* − *x*)^2^ / ∑(*y* − *ȳ*)^2^, where the sum extends to all species and replicates and *ȳ* is the mean of *y*. Formally, this value is related to the coefficient of determination of a 1:1 fit as 1 − *R*^2^. Therefore, inconsistencies greater than 1 are equivalent to *R*^2^ < 0, implying that the (*x*, *y*) pairs strongly deviate from a 1:1 trend (more precisely, the *y*-values are better predicted by their mean than by the *x*-values).

A robust estimate of the core genome size in mixed pangenomes was obtained with the command mOTUpan [42], using as inputs the OGC presence-absence matrices (option ‘-c’) and the completeness and contamination values reported in the GTDB and GMBC metadata files (option ‘-k’). Moreover, mOTUpan estimates of the mean genome size and the core genome size were used to derive an alternative measure of gene content diversity that accounts for genome incompleteness. Given a predicted core genome with size *C* and mean predicted genome size *G*, the gene content diversity measure is obtained as 1 − *C*/*G*. Assuming that the true genome sizes are similar among strains of the same species, this value is approximately equal to the proportion of genes within a genome that are accessory. Ready-to-use code to calculate the predicted percentage of accessory genes per genome from the output of mOTUpan is provided in Additional File 1.

## Declarations

## Ethics approval and consent to participate

Not applicable

## Consent for publication

Not applicable

## Availability of data and materials

The datasets generated and analyzed in this study are available in Zenodo, at https://dx.doi.org/10.5281/zenodo.7387758

## Authors’ contributions

JI and JHC designed the study; SMM and SGB generated the datasets for each clustering criterion; SMM, YL, SGB, and JI generated and compared the pangenomes; SMM, YL, JHC, and JI wrote the manuscript. All authors read and approved the final manuscript.

## Acknowledgements

We thank J. Botelho and L. Dudbridge for their feedback and critical read of the manuscript.

