## Supplementary Figures for "Comparison of gene clustering criteria reveals intrinsic uncertainty in pangenome analyses"

**Supplementary Figure S1**

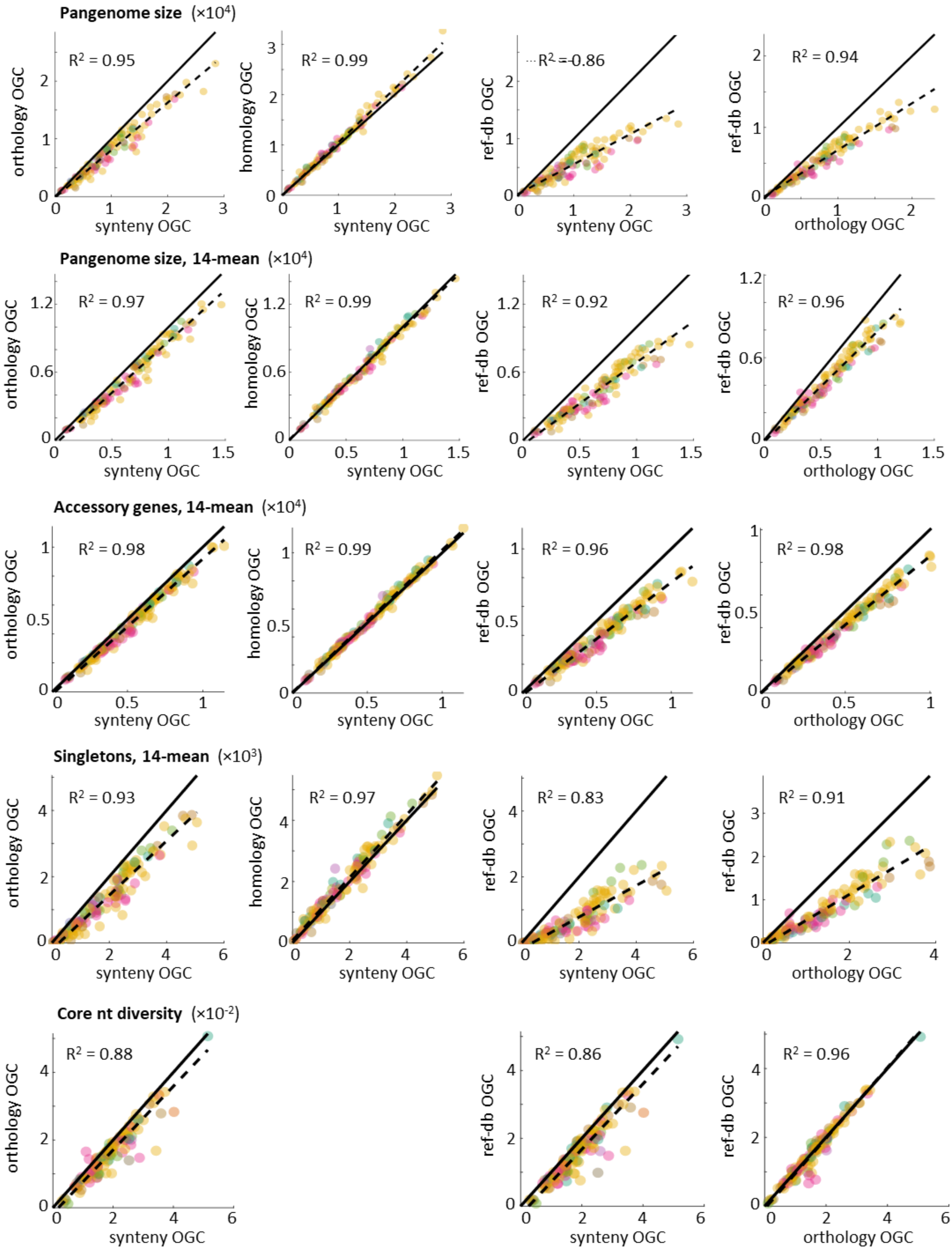


**Figure S1:** Species-wise comparison of pangenome features inferred from different sets of OGC. Dashed lines show the best orthogonal least-squares fit; solid lines indicate the 1:1 trend. Each point corresponds to the pangenome of one species, colored according to its phylum (see Figure 2 in the main text and Suppl. Fig. S3 for color-phyla mapping). For computational reasons, the core nucleotide diversity was not calculated for homology OGC.

**Supplementary Figure S2**

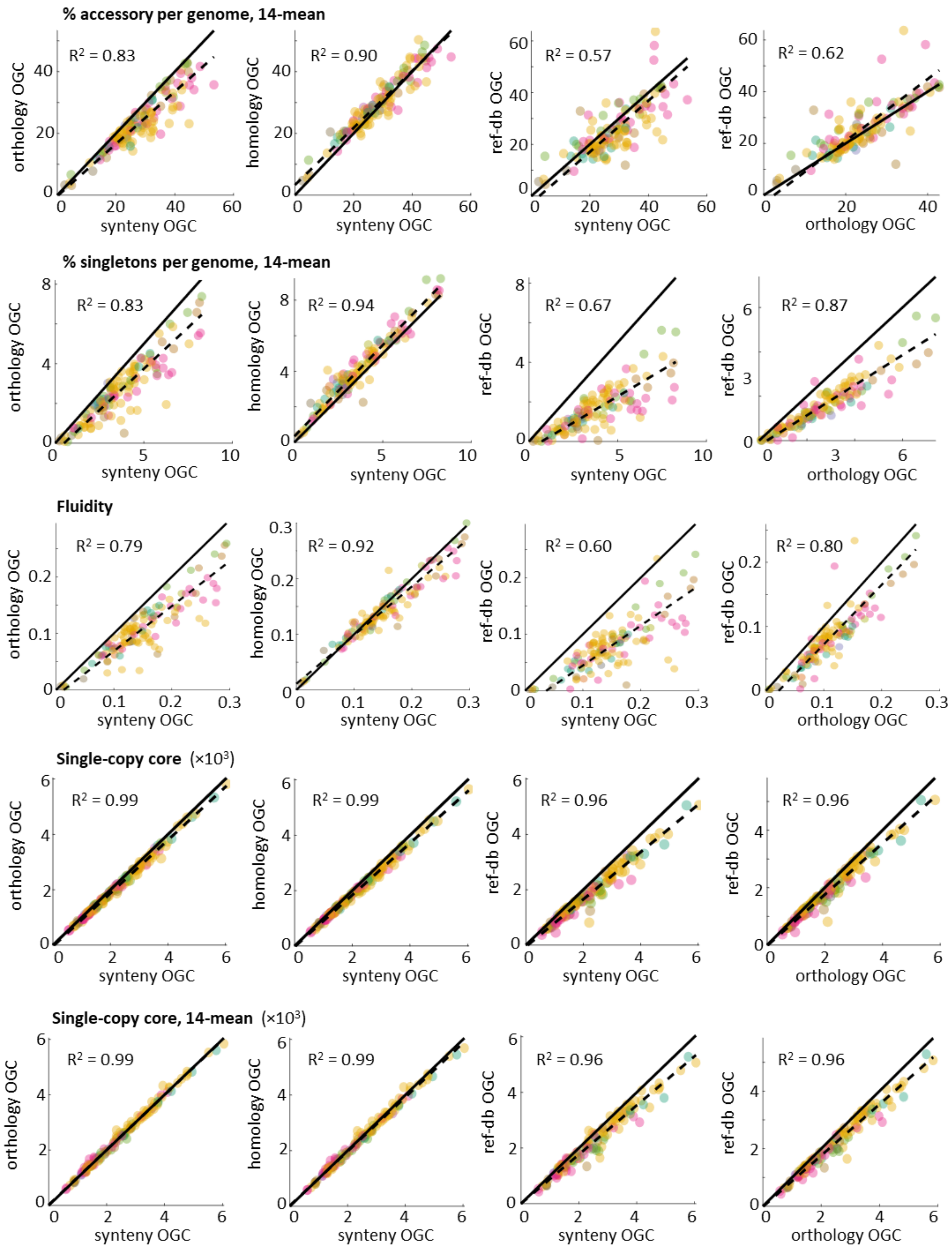


**Figure S2:** Species-wise comparison of pangenome features inferred from different sets of OGC. Dashed lines show the best orthogonal least-squares fit; solid lines indicate the 1:1 trend. Each point corresponds to the pangenome of one species, colored according to its phylum (see Figure 2 in the main text and Suppl. Fig. S3 for the color coding of phyla).

**Supplementary Figure S3**

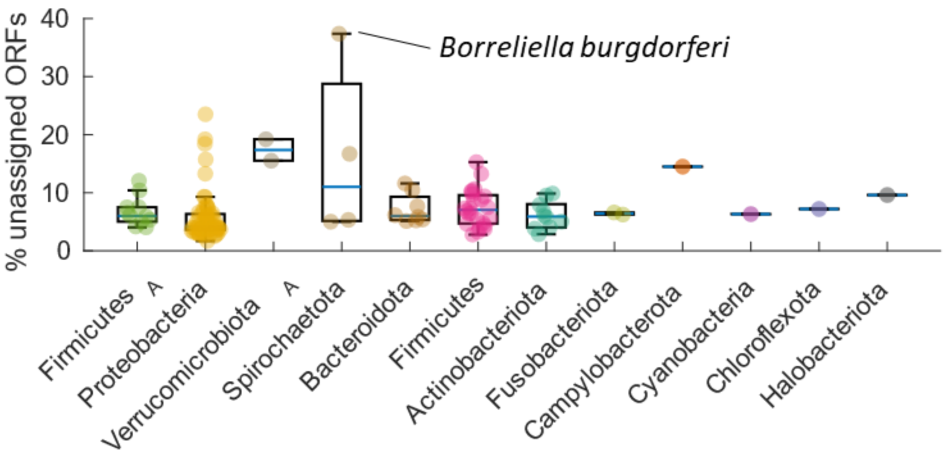


**Figure S3:** Fraction of ORFs per species that could not be classified into OGC by mapping to the eggNOG database. Species are grouped by phylum based on GTDB taxonomy.

**Supplementary Figure S4**

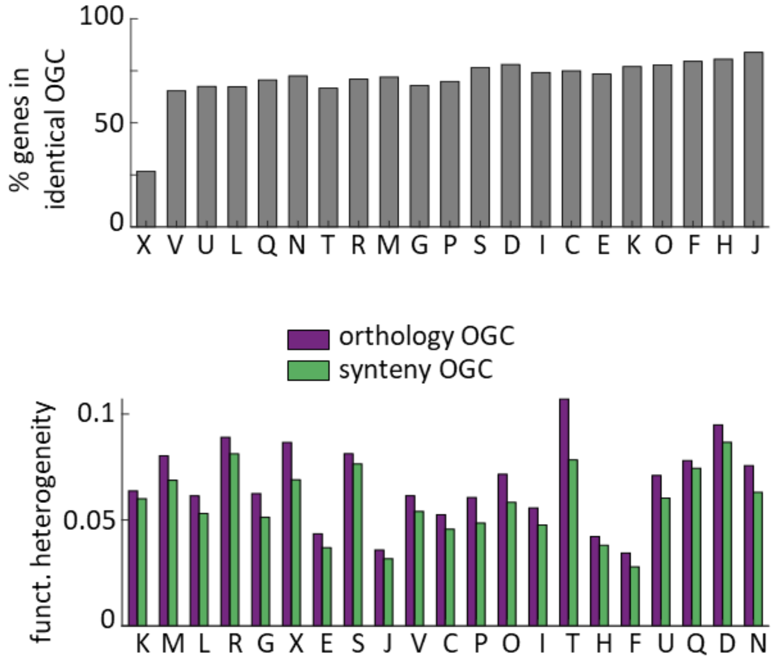


**Figure S4:** (a) Gene-level agreement between synteny- and orthology-based OGC, measured as the fraction of ORFs assigned to identical OGC and stratified by functional category. Identical OGC are those that contain exactly the same ORFs in both methods. (b) Functional heterogeneity in orthology- and synteny-based OGC. An OGC is considered functionally heterogeneous if it contains genes from >1 functional category (based on COG2020 functional annotation scheme). Abbreviations of functional categories, C: energy production and conversion; D: cell cycle control, cell division, chromosome partitioning; E: amino acid transport and metabolism; F: nucleotide transport and metabolism; G: carbohydrate transport and metabolism; H: coenzyme transport and metabolism; I: lipid transport and metabolism; J: translation, ribosomal structure and biogenesis; K: transcription; L: replication, recombination and repair; M: cell wall/membrane/envelope biogenesis; N: cell motility; O: posttranslational modification, protein turnover, chaperones; P: inorganic ion transport and metabolism; Q: secondary metabolites biosynthesis, transport and catabolism; R: general function prediction only; S: function unknown; T: signal transduction mechanisms; U: intracellular trafficking, secretion, and vesicular transport; V: defense mechanisms; X: mobilome.

### Supplementary Figure S5

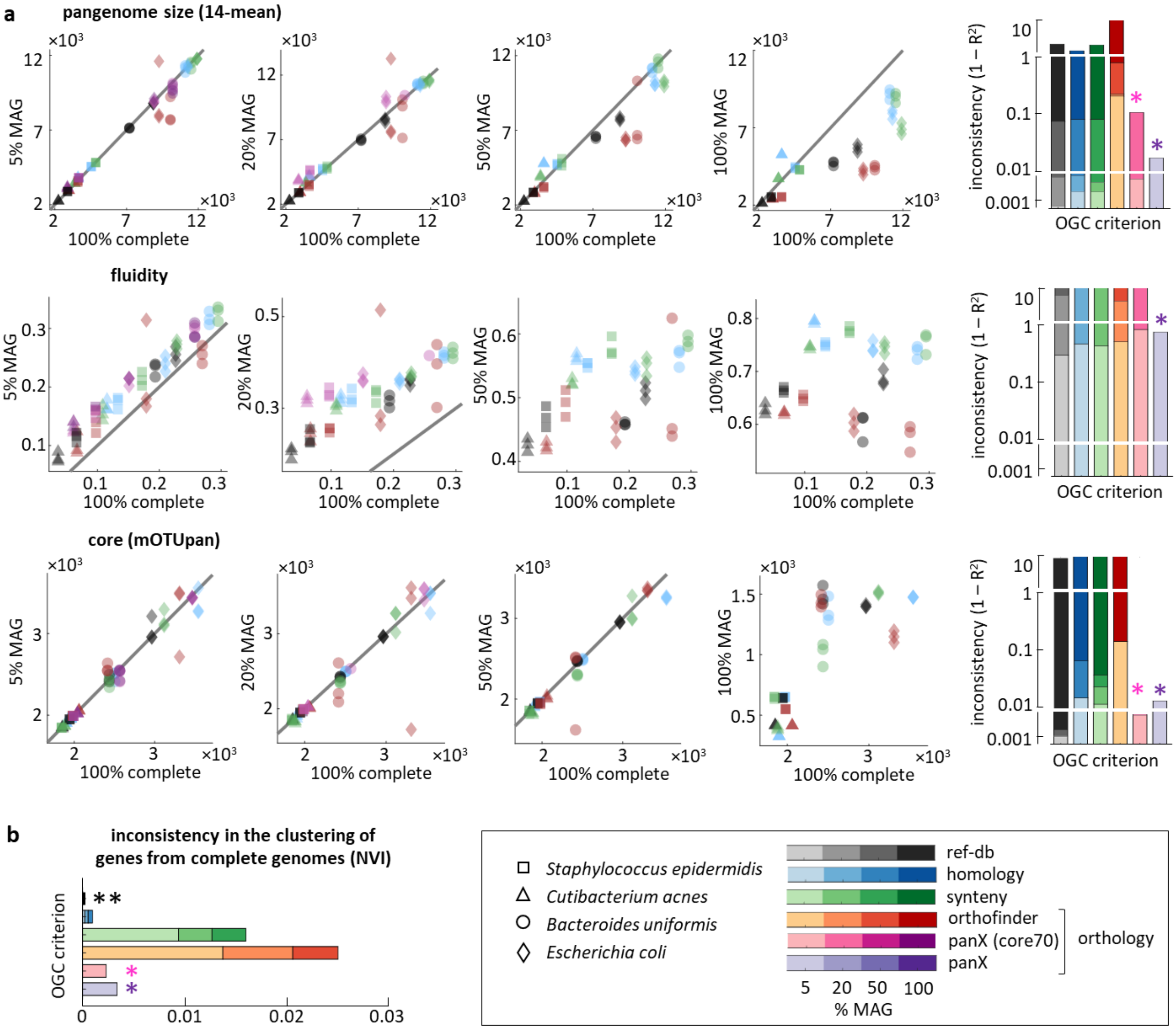


**Figure S5:** Comparison between pangenome properties inferred from high-quality genomes (x-axis) and mixtures of low-quality MAG and high-quality genomes (y-axis). From left to right, the scatter plots correspond to pangenomes with 5%, 20%, 50% and 100% of MAG. Each point in the scatter plots corresponds to one pangenome, with different symbols and colors used to distinguish among species and OGC generation methods, respectively. Each scatter plot combines data from 4 species, 6 gene clustering methods, and 3 random subsamples. The bar plots on the right summarize the observed inconsistencies, with color intensities representing the fraction of MAG. Note that panX fails to produce results in pangenomes that contain >5-10% of MAG (pink and purple asterisks). (b) Sensitivity of gene clustering methods to the addition of MAGs, calculated by comparing the cluster assignations of genes from high-quality genomes before and after adding MAG. NVI: normalized variation of information. Different OGC generation methods are color coded, with color intensities indicating the fraction of MAG. Note that reference-database mapping methods produce an NVI equal to zero (double asterisk).

**Supplementary Figure S6**

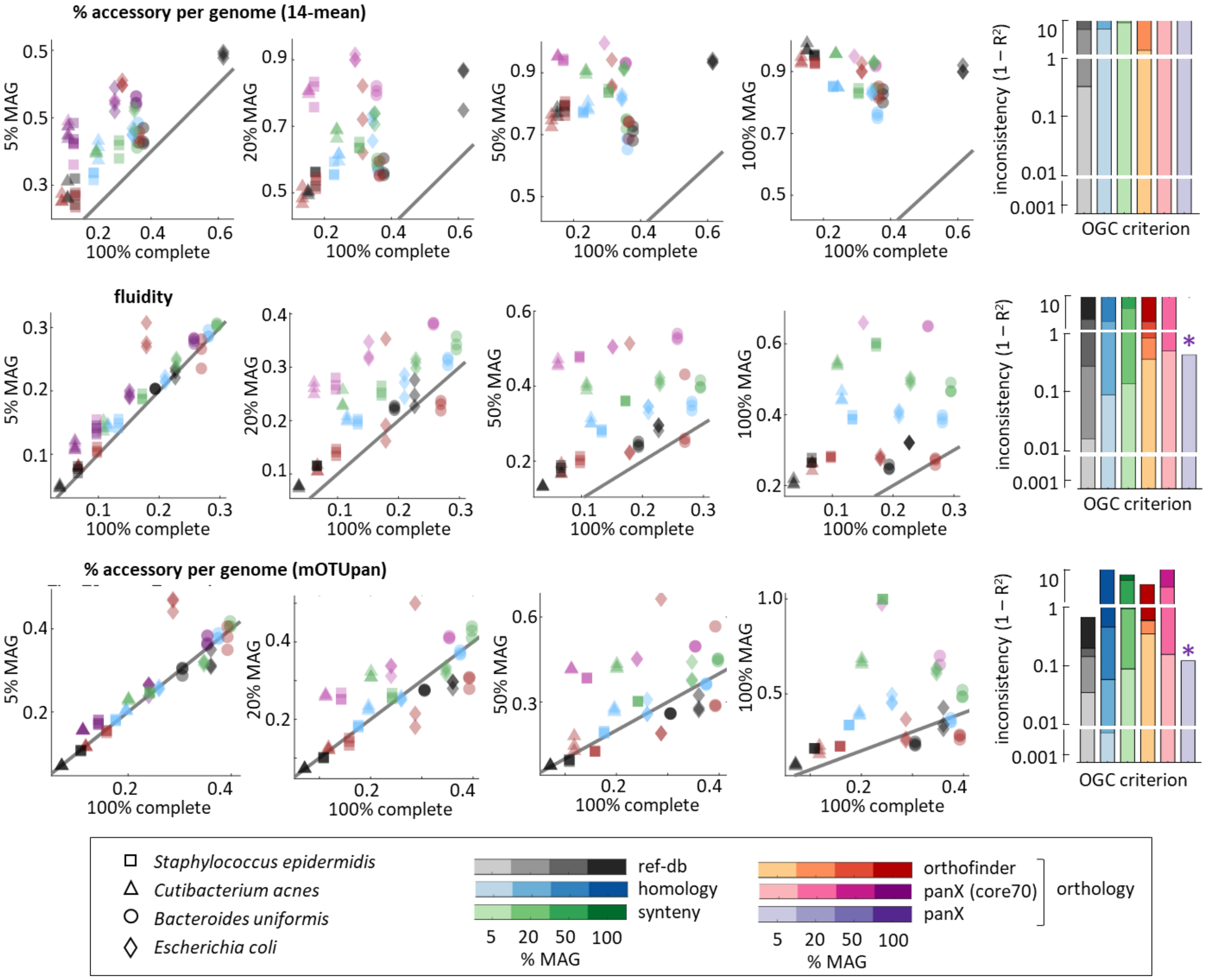


**Figure S6:** Consistency of different metrics of gene content diversity inferred from incomplete genomes. Each scatter plot compares the values obtained from high-quality genomes (x-axis) and mixtures of medium-quality MAG and high-quality genomes (y-axis). From left to right, the scatter plots correspond to pangenomes with 5%, 20%, 50% and 100% of MAG. Each point in the scatter plots corresponds to one pangenome, with different symbols and colors used to distinguish among species and OGC generation methods, respectively. Each scatter plot combines data from 4 species, 6 gene clustering methods, and 3 random subsamples. The bar plots on the right summarize the observed inconsistencies, with color intensities representing the fraction of MAG. Note that panX fails to produce results in pangenomes that contain >5% of MAG (purple asterisks). The panels for fluidity (middle row) are the same as those in Figure 6 and are reproduced here to facilitate comparison with other metrics.

**Supplementary Figure S7**

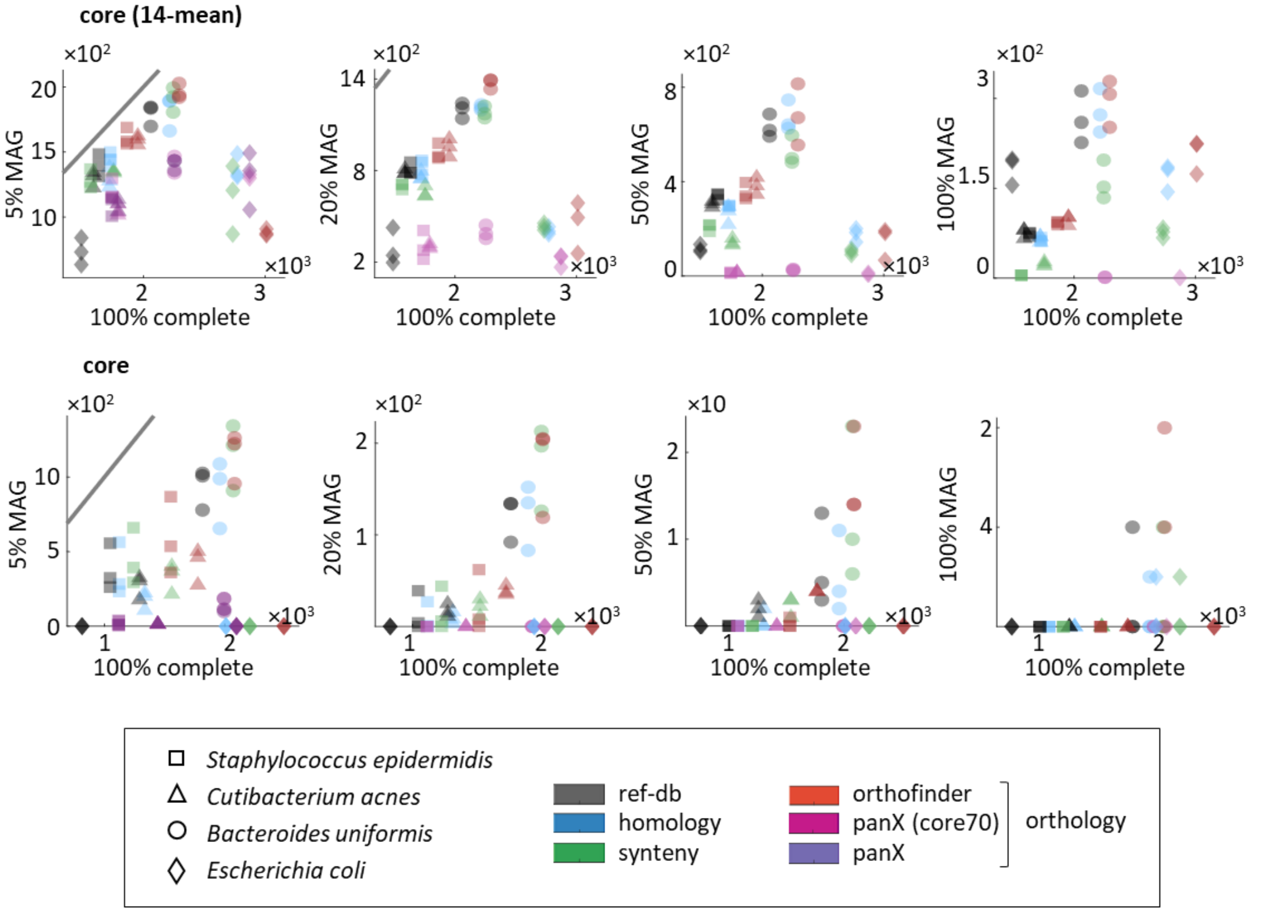


**Figure S7:** Consistency of different metrics of core genome size inferred from incomplete genomes. Each scatter plot compares the values obtained from high-quality genomes (x-axis) and mixtures of medium-quality MAG and high-quality genomes (y-axis). From left to right, the scatter plots correspond to pangenomes with 5%, 20%, 50% and 100% of MAG. Each point in the scatter plots corresponds to one pangenome, with different symbols and colors used to distinguish among species and OGC generation methods, respectively. Each scatter plot combines data from 4 species, 6 gene clustering methods, and 3 random subsamples.
