## Additional File 1 for "Comparison of gene clustering criteria reveals intrinsic uncertainty in pangenome analyses": PredictedAccessoryPerGenome.sh

```
#!/bin/bash
# Usage: ./PredictedAccessoryPerGenome.sh <mOTUpan_results_file>
# Or also: bash PredictedAccessoryPerGenome.sh <mOTUpan_results_file>
# Disclaimer: The script does not perform any sanity check. Please, check the input file before running it.
# Author: Jaime Iranzo (09/08/2023)
FILE=$1
echo "Results extracted from mOTUpan file: $FILE"
Core=$(grep '#core_length=' "$FILE" | cut -d'=' -f2) && echo "Estimated core genome size = $Core"
Gsz=$(grep '#mean_est_genome_size=' "$FILE" | cut -d'=' -f2 | cut -d'.' -f1) && echo "Mean estimated genome size = $Gsz"
PredAccPerGenome=$(bc <<< "scale=6 ; 1 - $Core / $Gsz") && echo "Predicted fraction of accessory genes per genome: $PredAccPerGenome"
```
